# Break-induced replication drives large-scale genomic amplifications in cancer cells

**DOI:** 10.1101/2024.08.27.609980

**Authors:** Mouadh Benamar, Rebeka Eki, Kang-Ping Du, Tarek Abbas

**Author notes:** Correspondence: Tarek Abbas, Ph.D.

## Abstract

DNA double-strand breaks (DSBs) are highly toxic lesions that underly the efficacy of ionizing radiation (IR) and a large number of cytotoxic chemotherapies^1–3^. Yet, abnormal repair of DSBs is associated with genomic instability and may contribute to cancer heterogeneity and tumour evolution. Here, we show that DSBs induced by IR, by DSB-inducing chemotherapeutics, or by the expression of a rare-cutting restriction endonuclease induce large-scale genomic amplification in human cancer cells. Importantly, the extent of DSB-induced genomic amplification (DIGA) in a panel of melanoma cell lines correlated with the degree of cytotoxicity elicited by IR, suggesting that DIGA contributes significantly to DSB-induced cancer cell lethality. DIGA, which is mediated through conservative DNA synthesis, does not require origin re-licensing, and is enhanced by the depletion or deletion of the methyltransferases SET8 and SUV4-20H1, which function sequentially to mono- and di-methylate histone H4 lysine 20 (H4K20) at DSBs to facilitate the recruitment of 53BP1-RIF1 and its downstream effector shieldin complex to DSBs to prevent hyper-resection^4–11^. Consistently, DIGA was enhanced in cells lacking 53BP1 or RIF1, or in cells that lacked components of the shieldin complex or of other factors that help recruit 53BP1 to DSBs. Mechanistically, DIGA requires MRE11/CtIP and EXO1, factors that promote resection and hyper-resection at DSBs, and is dependent on the catalytic activity of the RAD51 recombinase. Furthermore, deletion or depletion of POLD3, POLD4, or RAD52, proteins involved in break-induced replication (BIR), significantly inhibited DIGA, suggesting that DIGA is mediated through a RAD51-dependent BIR-like process. DIGA induction was maximal if the cells encountered DSBs in early and mid S-phase, whereas cells competent for homologous recombination (in late S and G2) exhibited less DIGA induction. We propose that unshielded, hyper-resected ends of DSBs may nucleate a replication-like intermediate that enables cytotoxic long-range genomic DNA amplification mediated through BIR.

## Main

Large-scale genomic amplification is a major source of genomic instability and may contribute to cancer cell cytotoxicity. We have previously shown that depletion of geminin (CDT1 inhibitor) or EMI1 (APC/C inhibitor) in head and neck squamous cell carcinoma (HNSCC) cells resulted in genomic amplifications resulting from re-replication, and this was stimulated by low doses of ionizing radiation (IR; 2-4 Gy)^12^. IR also stimulated re-replication in cells depleted, or acutely deleted, of CDT2, the substrate receptor of the CRL4^CDT2^ E3 ubiquitin ligase (leading to CDT1 and SET8 stabilization in S-phase), as well as in cells treated with the neddylation inhibitor MLN4924, which suppresses the activity of cullin-dependent E3 ligases, including CRL4^CDT2^ ^12^.

While studying the the response of cancer cells to IR ^12^, we observed a dose-dependent increase in the fraction of the head and neck squamouos carccinoma cells (HNSCC) FaDu or Cal27 cells (with mutant p53) that contained DNA content greater than 4N (Fig. 1a-c; highlighted in red), reminiscent of cells undergoing re-replication in response to MLN4924 treatment (Extended Data Fig. 1a). In these experiments, the cells were exposed IR, cultured for 48 hours, and labelled with propidium iodide (PI; to stain the DNA) and analysed by flow cytometry (FACS). The fraction of cells (Y axis) with DNA content of G1 (2N; absorbance at 200; X axis), G2/M (4N; absorbance at 400), or greater than 4N (>4N; absorbance >400; highlighted in red with frequency indicated on top) are blotted. This latter fraction (>4N) is often gated-out on FACS profiles, potentially explaining why such increase in aneuploidy/ploidy has not been fully characterized. The increase in genomic DNA following IR exposure was accompanied by an average expansion in nuclear size of ∼25%, reminiscent of cells undergoing re-replication (Fig. 1c). A similar increase in the fraction of cells with >4N DNA content was observed following the exposure of two additional p53-negative HNSCC cell lines to IR (Extended Data Fig. 1b, c). IR did not elicit significant genomic amplifications in benign hTERT-transformed OKF6-TERT keratinocytes (Fig. 1b). The induction of genomic amplifications following IR exposure was not limited to HNSCC cells, nor was it restricted to p53-negative cancer cells, as IR also induced large-scale genomic amplifications in human osteosarcoma U2OS cells (wild-type (WT) p53 – see below), in human colon cancer HCT116 (with WT p53) cells, in MDA-MB-231 breast cancer cells (mutant p53), and in HEK-293T cells (Fig. 1d, and Extended Data Fig. 1b-c). Deletion of *TP53* in the human colon cancer HCT116 (with WT p53) resulted in a larger increase in the fraction of cells exhibiting these amplifications following the exposure to various doses of IR, and this effect was more pronounced at lower IR doses (Fig. 1d), indicating that p53, which suppresses genomic amplifications (re-replication) that results from origin re-licensing^13^ (Extended Data Fig. 1d), also suppresses, but does not prevent, IR-induced genomic amplifications. As it will become evident below, we refer to this increase in DNA content as large-scale genomic amplification to distinguish it from aneuploidy or polyploidy that traditionally results from chromosomal mis-segregation or endoreduplication.

**Figure 1.**
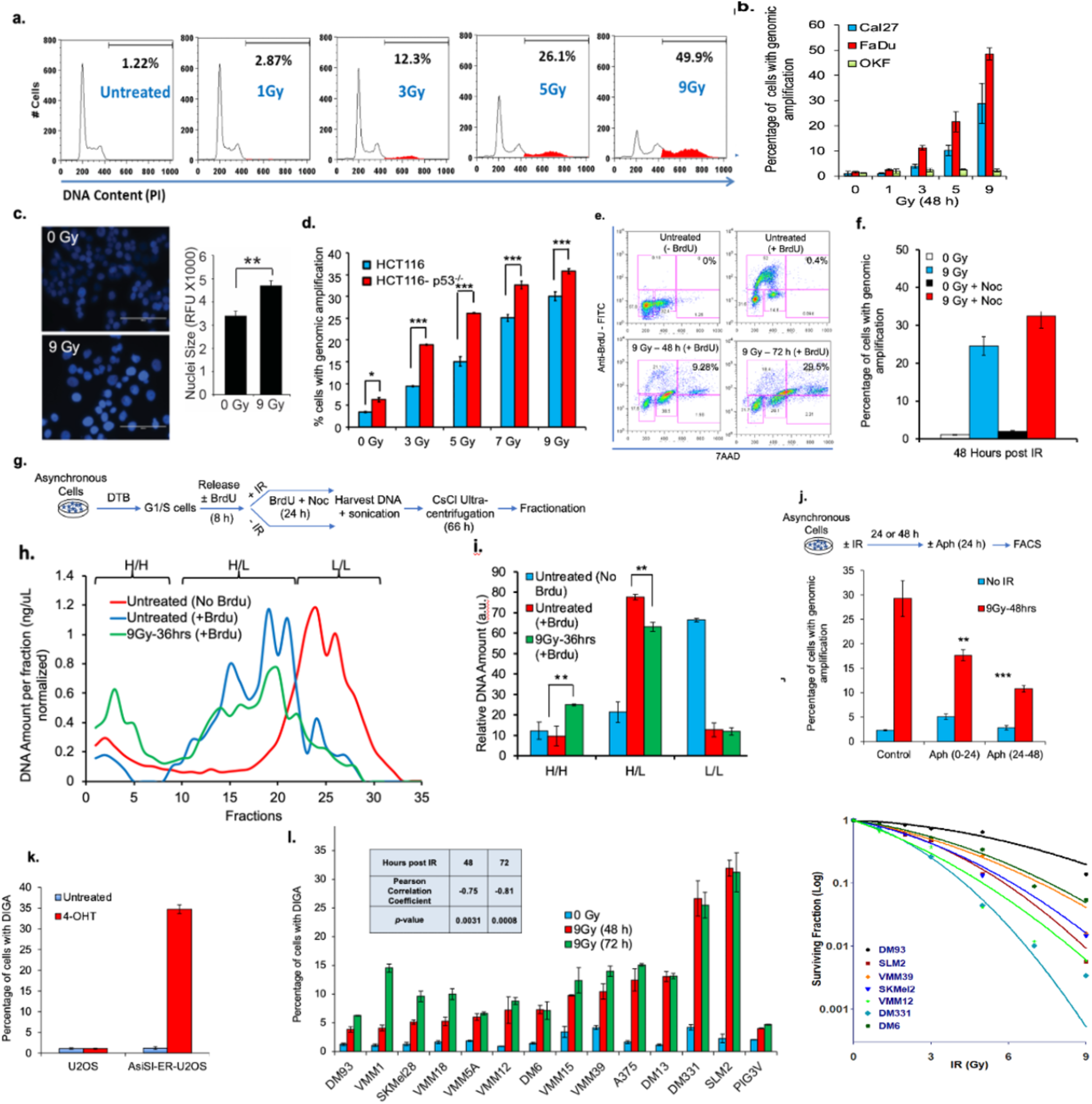
Induction of *bona fide* DNA genomic amplifications in cancer cells exposed to ionizing radiation or other DNA double strand break-inducing agents. **a,** Flow cytometry profile showing the increase in genomic content greater than G2/M (highlighted in red) in FaDu cells exposed to increasing doses of IR. **b**, Histogram showing the extent of genomic amplifications in Cal27 and FaDu HNSCC cell lines as well as the h-TERT immortalized keratinocytes (OKF) in response to increasing doses of IR. Genomic amplifications were examined 48 h post-IR. **c**, (*left*) Representative images of DAPI-stained Fadu nuclei following exposure to 9 Gy. (*right*) Quantitation of the nuclear size of FaDu cells 72 h following the exposure to 9 Gy. **d**, Genomic amplifications in HCT116 and p53-null (p53-/-) cells at various doses of IR measured 72 h post-IR. **e**, BrdU-7AAD FACS profiles of FaDu cells exposed to 9 Gy and pulse-labelled with BrdU for 1 h, 48 h post-IR. **f**, Genomic amplifications in U2OS cells exposed to 9 Gy, and treated with or without nocodazole at 24 h post-IR. Genomic amplifications were examined 48 h post-IR. **g,** Experimental workflow of the various treatments of U2OS cells for the detection of IR-induced genomic amplifications by DNA isotope labelling and CsCl ultracentrifugation and fractionation. **h**, Plot showing the DNA content of each CsCl fraction in cells treated according to the workflow in **g**. **i.** Histogram showing the relative amount of DNA detected in the fractions corresponding to non-replicating (L:L), singly replicated (H:L), and doubly replicated (H:H; doubly replicated) DNA as determined by measuring the area under the curve (shown in **h**) from three independent experiments (n=3). **j**, Histogram showing the percentage of U2OS cells with rereplication 48 h following the exposure 9 Gy with or without aphidicolin (Aph) treatment according to the schematic shown on top. **k**, The percentage of U2OS cells with genomic amplifications following the expression of A*si*SI restriction endonuclease by 4-OHT treatment for 72 h. **l**, Histogram showing the percentage of the indicated melanoma cell lines with genomic amplifications measured at 48 and 72 h following exposure to 9 Gy (n=3). Correlation and statistical significance of the induction of genomic amplifications both at 48 and 72 h post-IR and IR-induced cytotoxicity in all the melanoma lines shown, defined as surviving fraction at 10% of the parental population (SF10), is shown on the top-right corner. Right: Representative clonogenic survival curves of a subset of the melanoma lines used. Data in all figures represent the average of three independent experiments ± S.D. *p*-values were calculated using Student’s *t*-test: \**p* < 0.05, \*\**p* < 0.01, \*\*\**p* < 0.001.

### IR-induced genomic amplifications involve newly-synthesized genomic sequences and does not require cells’ entry into mitosis

To begin to characterize the mechanism underlying the genomic amplifications by IR, we focused on U2OS cells, a commonly used cancer cell line used in these studies. Genomic amplifications in U2OS cells are readily detectable by FACS as early as 15 hrs post-IR (∼8%), but reaches significant levels at 36 (∼20%) and 48 hrs (∼35%) post-exposure (Extended Data Fig. 2a, b). The bulk of DNA amplifications were between 5N and 6N DNA content, but with a significant fraction of cells accumulating 6-8N and >8N DNA content, both of which increased with increasing radiation dose (Extended Data Fig. c-e). These large increases in genomic DNA content following IR exposure, therefore, cannot possibly be attributed to insertional mutagenesis, which usually incorporate only few nucleotides at DSBs, especially that most DSBs are repaired through processes that result mostly in deletions and/or translocations, rather than insertions. We reasoned these amplifications may result from: (a) chromosomal mis-segregation or failed cytokinesis, (b) mitosis bypass or endoreduplication, (c) some form of genome-wide re-replication, or (d) large-scale, repair-associated DNA synthesis. To distinguish between these possibilities, we first exposed U2OS cells to 9Gy, and, 48 hrs post-IR, the cells were pulse-labelled with bromodeoxyuridine (BrdU) for 1 hr to visualize active DNA synthesis. Although the S-phase population was significantly depleted by this treatment, more than 25% of the irradiated cells contained >4N DNA content that stained positive for BrdU (Fig. 1e). The bulk of the DNA synthesis occurred in cells with >4N, but with <8N DNA content, indicating that DNA synthesis occurs within the same cell division cycle. Similar results were found in U2OS cells exposed to IR and pulse-labelled with BrdU (Extended Fig. 3a). These results indicate that either cells with mis-segregated DNA entered a second round of S phase or that irradiated cells never entered mitosis in the first place, and underwent a new round of DNA synthesis in late S or in G2. In support of the latter possibility, treatment of U2OS with nocodazole, which arrest the cells in prometaphase, did not reduce the fraction of IR-treated cells with >G2/M DNA content (Fig. 1f, and Extended Data Fig. 3b). Thus, the observed increase in genomic content in IR-treated cells appeared to result from new round(s) of DNA synthesis that occurred within a single cell division cycle, and additionally, does not require the entry of cells into mitosis.

To substantiate this conclusion, we utilized caesium chloride density gradient centrifugation of genomic DNA to enable the separation of doubly-labelled DNA (heavy-heavy; H:H) from singly replicated (heavy-light; H:L) DNA following BrdU labelling ^14^. To overcome the immediate general inhibition of DNA synthesis that follows IR exposure and to permit the labelling of H:L DNA, U2OS cells were arrested at G1/S with double thymidine block (DTB), and subsequently released into S-phase in the presence of BrdU (Fig. 1g). Eight hours later, the cells were exposed to 9 Gy and treated with nocodazole to prevent slippage into the next cell cycle, and DNA was purified 36 hrs post-IR. Whereas the mock-treated cells contained mostly H:L DNA, a significant fraction of H:H DNA was readily observed in irradiated cells (Fig. 1h, i, and Extended Data Fig. 3c). To test whether the increase in genomic DNA content induced by IR was mediated by replicative polymerases operating during S phase (POLα, ο and χ), we exposed U2OS cells to 9 Gy, and treated the cells with aphidicolin (2 µM) either immediately following radiation or 24 hrs post-IR; cells were harvested 48 hrs post-IR. In both cases, aphidicolin treatment suppressed genomic amplifications induced by IR (Fig. 1j, and Extended Data Fig. 3d). The accumulation of H:H DNA extracted from cells released from the DTB and exposed to IR was similarly suppressed if the cells were treated with aphidicolin along with nocodazole (Extended Data Fig. 3e). This result indicates that genomic amplifications result from at least two rounds of new semiconservative DNA synthesis within the same cell cycle (i.e., re-replication) or from some form of repair-associated conservative DNA synthesis, similar to that employed by break-induced replication (BIR), a homologous recombination pathway used for the repair of collapsed replication forks ^15–19^.

IR is a serious environmental catastrophe for a cell; it induces a variety of DNA lesions in addition to DSBs including single strand breaks, base lesions, apurinic/apyrimidinic sites (abasic sites), sugar damage, and clustered DNA lesions^20,21^. We asked whether DSBs elicited by any means other than IR induced genomic amplifications. We first tested whether agents with the capacity to induce DSBs will induce amplifications similar to IR. We found that treatment of U2OS cells with the topoisomerase II inhibitor, etoposide (1 µg/ml), caused significant amplifications (Extended Data Fig. 3f). The treatment of U2OS with low dose doxorubicin (0.1 µM), which induces DSBs primarily during DNA replication, also induced genomic amplifications, but higher concentrations resulted in robust apoptosis that precluded the accurate measurement of these amplifications (Extended Data Fig. 3f, and data not shown). On the other hand, exposure of U2OS to ultraviolet radiation (100 J/m^2^), which mostly cause bulky DNA adducts but few DSBs, resulted in minimal genomic amplifications (Extended Data Fig. 3f). These results suggest that large-scale genomic amplifications results primarily from DSBs, but not from other assaults that disturb the integrity of the DNA backbone or damage its adducts. We next asked whether DSBs are sufficient to induce genomic amplifications in cancer cells. DSBs were induced in U2OS cells via the endogenous expression of the *Asi*SI restriction enzyme gene fused to a gene encoding the oestrogen receptor (*Asi*SI-ER-U2OS cells^22^). *Asi*SI is an 8-bp cutter that, in practice, delivers ∼150 DSBs per human genome^22–24^. Treatment with 300 nM 4-OHT induced aphidicolin-sensitive increase in genomic DNA content in 35% of *Asi*SI-ER-U2OS cells, but had no effect on normal U2OS cells (Fig. 1k, and Extended Data Fig. 3g). Furthermore, pulse labelling of *Asi*SI-ER-U2OS cells with BrdU at various hrs following 4-OHT treatment further confirmed that genomic amplifications occur within the same cell division cycle (Extended Data Fig. 3h). Because genomic amplifications appear to result from DSBs, but not other assaults that disturb the integrity of the DNA backbone or damage its adducts, we refer to these large-scale genomic amplifications as DSB-induced genomic amplifications (DIGA).

Previous studies have shown that cells with structural or numerical chromosomal instability are more susceptible to radiation exposure. Because DIGA is associated with increased genomic DNA content, we hypothesized that cells that exhibiting increased DIGA in response to radiation exposure will be more effectively suppressed than those that undergo limited or low levels DIGA. To test this hypothesis, we examined the susceptibility of panel of thirteen melanoma cell lines to the induction of genomic amplifications following the exposure to 9 Gy and monitored DIGA induction at 48 and 72 hrs post-radiation by FACS. As a control, we also exposed the non-malignant PIG3V melanocytes to the same treatment. While less than 5% PIG3V cells exhibited DIGA following IR treatment, the majority of melanoma lines tested exhibited significant but varying degrees of DIGA, that, in most cases, increased with time (Fig.1l). These melanoma cell lines exhibited varying degree of proliferation rates, which modestly correlated with the extent to which these various lines exhibited DIGA at 48 (R^2^=0.2661) or 72 (R^2^=0.1856) hrs post-irradiation (Extended Data Fig. 4a-c). We next performed clonogenic survival assays for these melanoma lines by exposing them to increasing IR doses, and monitoring their ability to grow colonies two weeks following exposure. From this analysis, we found that the propensity of the various melanoma lines to induce DIGA, both at 48 and 72 h post-exposure, strongly correlated (R^2^ = 0.5624 and 0.6492, respectively) with IR-induced toxicity (Figures 1l, and Extended Data Fig. 4d, e). Thus, DIGA appears to contribute significantly to IR-induced cytotoxicity.

We next sought to identify the mechanism by which DIGA is induced in cancer cells using IR as the source of DSB induction. The extent of genomic amplifications induced by DSBs indicate that the “extra” DNA must arise from illicit template amplification either via reactivation of *bona fide* replication origins or by an extensive form of DNA repair synthesis. The cullin 4 E3 ubiquitin ligase complex CRL4^CDT2^ is a major suppressor of origin re-firing and genome-wide re-replication induction in cancer cells^25,26^. CRL4^CDT2^ suppresses re-replication primarily through its ability to promote the ubiquitylation and degradation of the replication licensing factors CDT1 in S and G2 phases of the cell cycle (Fig. 2a). Indeed, the FACS profiles of DIGA induction in U2OS cells exposed to 9 Gy were similar to the re-replication FACS profiles of cells with reactivated replication origins resulting from treatment with the neddylation inhibitor MLN4924^12,27,28^, which inhibits all cullin-based E3 ligases including CRL4^CDT2^ (Extended Data Fig. 1a). DNA damage such as that induced by IR, however, provokes CRL4^CDT2^ activation, resulting in the degradation of CDT1^27,29–31^. To test whether DIGA results from failure to degrade CDT1, potentially through defects in CRL4^CDT2^ activity, we exposed Cal27 cells to 4 Gy and monitored CDT1 levels by immunoblotting (Fig. 2b). The results show that CDT1 was efficiently degraded following IR exposure in these cells. Importantly, the depletion of CDT1 by siRNA prior to IR exposure had no impact on DIGA induction, although it readily suppressed re-replication induction by MLN4924 (Fig. 2c, and Extended Data Fig. 5a). Thus, DIGA observed following IR exposure is not a consequence of canonical CDT1-dependent origin re-licensing.

**Figure 2.**
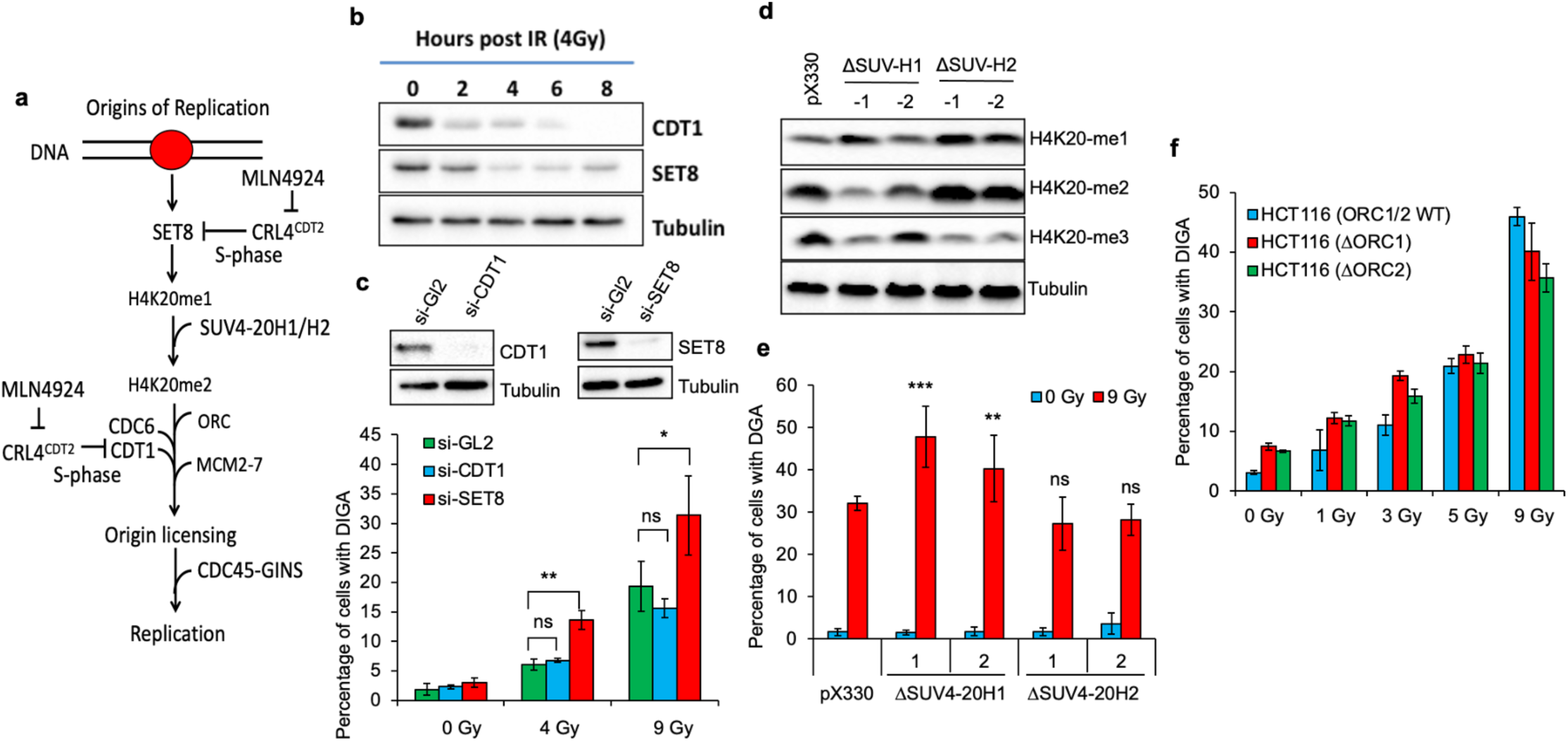
DIGA is not mediated by ORC1-2/CDT1-dependent origin licensing, and is suppressed by SET8/SUV4-20H1. **a**, Schematic of the various molecules involved in origin licensing in mammalian cells in which SET8 and CDT1 are targeted for degradation by CRL4^CDT2^ during S-phase to suppress origin re-licensing. **b**, Immunoblots of CDT1 and SET8 in Cal27 cells exposed to IR (4 Gy) and analysed at the indicated time points post-IR (4 Gy). **c,** (*top*) immunoblotting of CDT1 and SET8 in Cal27 following treatment with the indicated siRNAs. (*bottom*) Histogram showing the percentage of control (siGL2) Cal27 cells or Cal27 cells with genomic amplifications following the depletion of CDT1 or SET8 by siRNA (for 48 h) and subsequent exposure to 4 or 9 Gy (analyzed 48 hr post irradiation). **d**, Immunoblot showing the impact of CRISPR/Cas9-mediated deletion of SUV4-20H1 or SUV4-20H2 in U2OS cells on histone H4K20 methylation. **e**, Histogram showing the percentage of U2OS cells undergoing genomic amplifications following the deletion of SUV4-20H1 or SUV4-20H2 (two independent clones each are shown). **f**, The impact of ORC1 or ORC2 deletion in HCT116 on IR-induced genomic amplifications. The histogram shows the percentage of control HCT116 or HCT116 deleted of ORC1 (1ORC1) or ORC2 (1ORC2) undergoing genomic amplifications 72 h following the exposure to the indicated doses of IR. Data in the histograms represent the average of three independent experiments ± S.D. ns: non-significant, \**p* < 0.05, \*\**p* < 0.01, \*\*\**p* < 0.001.

Origin re-licensing and re-replication induction are also triggered by the failure to promote CRL4^CDT2^-dependent degradation of SET8 during S phase^28,32–42^ (Fig. 2a). *SET8* is an essential gene that encodes a histone mono-methyltransferase for H4K20^43,44^. The SET8-dependent re-replication requires the conversion of H4K20me1 to H4K20me2 by SUV4-20H1/H2; this helps recruit the origin recognition complex (ORC) through the BAH domain of ORC1, which recognizes and binds H4K20me2 at replication origins^45–47^. However, exposure of Cal27 to 4 Gy resulted in a time-dependent degradation of SET8 protein, albeit with slower kinetics than that observed for CDT1 (Fig. 2b). Furthermore, when SET8 was depleted by siRNA prior to IR exposure, which would be expected to suppress DIGA if an SET8-mediated and origin-dependent pathway was in play, the percentage of cells exhibiting DIGA was *increased* (Fig. 2c). DIGA was also stimulated in U2OS cells in which *KMT5B* (encoding SUV4-20H1) was deleted, but not in cells in which *KMT5C* (encoding SUV4-20H2) was deleted (Fig. 2d, e). Stimulation of DIGA induction following the deletion of *KMT5B* was confirmed in VMM39 melanoma cells, which do not normally undergo robust DIGA (Fig. 1l, and Extended Data Fig. 5b). Given that the deletion of *KMT5B*, but not *KMT5C,* reduced H4K20 dimethylation (Fig. 2d), we conclude that dimethylation of H4K20, catalysed by SUV4-20H1, is particularly important for suppressing DIGA. Importantly, when we irradiated HCT116 cells that had sustained deletions of *ORC1*, which are essential for mediating the re-replication induction by SET8{Ref}, or *ORC2*^48^, DIGA was induced to levels comparable to that observed in parental HCT116 cells with WT ORC1/2 (Fig. 2f). Thus, while DNA replication in these ORC1/2-deficient cells may be mediated via other ORC proteins or through a bypass mechanism that requires the CDC6 replication licensing protein^48^, our data indicate that DIGA cannot be mediated by SET8- or H4K20me-dependent recruitment of ORC1/2. Together, these results indicate that the mechanism of DIGA does not require SET8-ORC1-2/CDT1-dependent origin licensing, and is suppressed by a SET8- and SUV4-20H1-dependent mechanism.

In addition to their role in promoting origin licensing in late mitosis and early G1, SET8 and SUV4-20H1 are known to localize to DSBs, and to promote repair of DSBs by non-homologous end-joining (NHEJ) through the recruitment of the NHEJ protein 53BP1^4–6^. Thus, we hypothesized that the increased frequency of DIGA we see in cells depleted of SET8 or lacking SUV4-20H1 results from defective NHEJ repair. We generated isogenic U2OS cell lines lacking proteins directly involved in the repair of DSBs via the NHEJ pathway. We first focused on DNA-PKcs, the catalytic subunit of DNA-PK, since this kinase is not only important for promoting NHEJ through the canonical pathway (c-NHEJ), but also because it cooperates with KU70 to recruit SET8 to DSBs^6,30,49^. We found that DIGA was significantly stimulated in U2OS or 293T cells deleted of *PRKDC* (encoding DNAPKcs) (Fig. 3a, and Extended Data Fig. 6a). DIGA was also enhanced following the treatment of U2OS or 293T cells with the DNA-PKcs specific inhibitor NU7441 (Fig. 3b and Extended Data Fig. 6b), indicating that the catalytic activity of DNA-PKcs is essential for suppressing DIGA. The requirement for DNA-PKcs catalytic activity in suppressing DIGA was further confirmed in DM93 melanoma cells, which undergo limited genomic amplifications in response to IR (Fig. 1l and Extended Data Fig. 6c). Surprisingly, DIGA was significantly *suppressed* by the deletion of *XRCC4* or *NHEJ1* (encoding XRCC4-like factor, XLF), and to a lesser extent by deletion of *LIG4*, all of which are core components of c-NHEJ functioning downstream of DNA-PKcs in the c-NHEJ pathway (Fig. 3c, d). Inhibition of DNA LIG4 activity also suppressed DIGA (Fig. 3e). The suppression of DIGA in cells lacking effectors of c-NHEJ was not a general requirement for these factors in resolving re-replication intermediates since cells lacking the genes coding these proteins re-replicated their DNA to the same extent as control U2OS cells following MLN4924 treatment (Extended Data Fig. 6d). Because we see more a dramatic suppression of DIGA following the deletion of XRCC4/XLF than that seen following the deletion or inhibition of LIG4, the results suggest that filament formation and stabilization of the DSBs broken ends by the LIG4/XRCC4/XLF complex^50,51^, rather than the eventual ligation of broken ends, is critical for DIGA induction. Thus, although the effectors of c-NHEJ appear to be required for DIGA induction, upstream components of this pathway (SET8, SUV4-20H1, and DNA-PKcs) all suppress this form of genomic amplifications. DIGA was not impacted by the deletion of *LIG1*, which is important for DSB repair effected by single-strand annealing (SSA)^52^ (Extended Data Fig. 6e). On the other hand, deletion of *POLQ,* an mediator of repair of DSBs via the microhomology-dependent alternative branch of NHEJ (alt-NHEJ), and a known suppressor of DSB repair by homologous recombination (HR)^53^, stimulated DIGA (Extended Data Fig. 6f).

**Figure 3.**
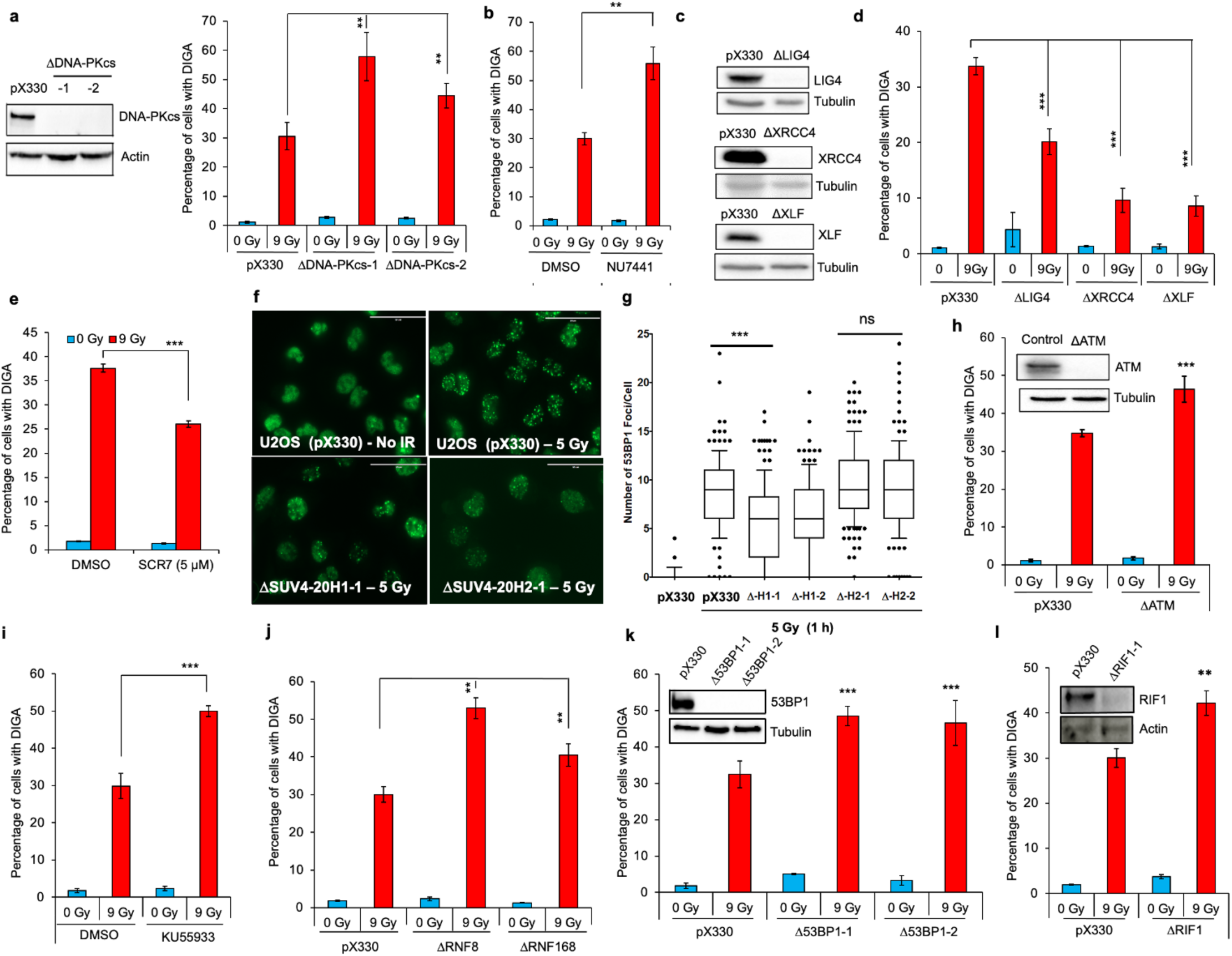
The impact of various proteins involved in the repair of DSBs on IR-induced genomic amplifications. **a**, Histogram (*right*) showing the impact of increasing IR doses on the induction of genomic amplifications in control U2OS cells or two independent clones with deletion of DNA-PKcs by CRISPR/Cas9. (*left*) Western blots illustrating the efficiency of deletion of DNA-PKcs in U2OS cells. **b**, Histogram showing the percentage of U2OS cells with genomic amplifications 72 h following the exposure to 9 Gy of control, DMSO-treated U2OS cells or U2OS cells treated with the DNA-PKcs inhibitor NU7441 for 24 h prior to IR. **c,** Immunoblot showing deletion of effectors of the c-NHEJ pathway (LIG4, XRCC4, XLF). **d**, Histogram showing the impact of deletion of the indicated effectors of c-NHEJ on the induction of genomic amplifications by IR. **e**, The impact of inhibiting DNA LIG4 on the induction of genomic amplifications by IR in U2OS cells. **f**, **g**, Deletion of *KTM4B* (coding for SUV4-20H1) but not deletion of *KTM4C* (coding for SUV4-20H2) inhibits the recruitment of 53BP1 to DSBs. **f**, Representative immunofluorescent images of 53BP1 showing formation of 53BP1 foci following exposure of control (pX330) U2OS cells or U2OS cells deleted of *KTM4B* or *KTM4C* to 5 Gy analysed 1 h post-exposure. **g**, Quantitation of the number of 53BP1 foci per cell in control (pX330) cells or U2OS deleted of *KTM4B* or *KTM4C* (two independent clones each). The results represent the average of a minimum of 100 nuclei in each of three independent experiments for each condition ± S.D. ns: non-significant, \*\*\**p* < 0.001. **h**, **i**, Histograms showing the percentage of cells undergoing genomic amplifications following the exposure of control or ATM-deleted U2OS cells (inset: immunoblotting of ATM levels) to 9 Gy (**h**), or following pharmacological inhibition of ATM in parental U2OS cells by KU55933 24 h prior to IR (**i**). Genomic amplifications was monitored 72 h post-IR. **j**, Histogram showing the percentage of control (pX330) U2OS cells or U2OS cells deleted of RNF8 or RNF168 and exposed to 9 Gy. Genomic amplifications was monitored 72 h post-IR. **k**, **l**, Histograms showing the percentage of control (pX330) U2OS cells or U2OS cells deleted of 53BP1 (in two independent clones; immunoblots shown in inset (**k**)) or RIF1 (immunoblots shown in inset (**l**)) with genomic amplifications following the exposure to 9 Gy. Genomic amplifications was monitored 72 h post-IR. Data in all the histograms represent the average of three independent experiments ± S.D. \**p* < 0.05, \*\**p* < 0.01, \*\*\**p* < 0.001.

The SET8-dependent mono-methylation of H4K20 at DSBs is converted to H4K20me2 by SUV4-20H1/H2, which is essential, albeit not sufficient, for recruiting the DSB repair protein 53BP1 to DSBs to antagonize HR and promote NHEJ^4–6^. We found that, similar to their differential impact on IR-induced DIGA (Fig. 2e, f), IR-induced 53BP1 foci formation was significantly diminished in U2OS cells deficient in SUV4-20H1 but not SUV4-20H2 (Fig. 3f, g). To test whether 53BP1 was critical for suppressing DIGA independent of H4K20 methylation, we deleted, in U2OS cells, several genes that are important for 53BP1 recruitment to DSBs, including the ataxia telangiectasia mutated kinase (ATM), the mediator of DNA damage checkpoint proteins 1 (MDC1), and the ubiquitin ligases RNF8 and RNF168^54^. We found that deletion of *ATM* or its pharmacological inhibition by the ATM specific inhibitor KU55933 stimulated DIGA induction by IR (Fig. 3h, i). ATM inhibition also robustly stimulated DIGA in A*si*SI-ER-U2OS following tamoxifen treatment (Extended Data Fig. 7a), but had no effect on MLN4924-induced re-replication in normal U2OS cells (Extended Data Fig. 7b). DIGA induction by IR was similarly stimulated by the deletion of *MDC1*, *RNF8*, or *RNF168*, each of which, as expected, exhibited deficiencies in 53BP1 recruitment to DSBs (Fig. 3j and Extended Data Fig. 7d-e). To examine if failure to recruit 53BP1 to DSBs impacted DIGA directly, we deleted *TP53BP1* (encoding 53BP1) in U2OS (and in 293T) cells by CRISPR/Cas9, and found that IR-induced DIGA was stimulated in these cells (Fig. 3k, and Extended Data Fig. 8a). Furthermore, deletion of *TP53BP1* in VMM39 melanoma cells, which undergo only modest genomic amplifications following IR exposure also stimulated DIGA induction (Fig. 1l and Extended Data Fig. 8b). 53BP1 deletion, on the other hand, did not impact re-replication induction by MLN4924 (Extended Data Fig. 8c). Furthermore, stimulation of DIGA was also observed in cells lacking the 53BP1 cofactor RIF1, whose deletion modestly inhibited the recruitment of 53BP1 to DSBs (Fig. 3l, and Extended Data Fig. 7c, d).

The main function of 53BP1-RIF1 at DSB is to protect against hyper-resection of the broken DNA ends through the recruitment of the shieldin complex^7–11^. Thus, we hypothesized that hyper-resection at DSBs plays a key role in DIGA induction. Consistent with this hypothesis, we found that silencing the expression of CtIP or MRE11, which initiate end-resection at DSBs, or the exonuclease EXO1, which promotes hyper-resection in U2OS significantly diminished DIGA (Fig. 4a, b and data not shown). Furthermore, deletion of SHLD1, SHLD2, or SHLD3, components of the shieldin complex, stimulated DIGA without compromising 53BP1 recruitment to DSBs (Fig. 4c, d and Extended Data Fig. 9a, b). Importantly, silencing or pharmacological inhibition of RAD51 in U2OS cells inhibited DIGA induction in a dose-dependent manner (Fig. 4e, f and Extended Data Fig. 9c), indicating that strand invasion by the hyper-resected DSBs is crucial for DIGA initiation. Finally, we found that DIGA was stimulated in U2OS cells depleted of factors involved in break-induced replication, including RAD52, POLD3 and POLD4 (Fig. 4g-i), consistent with the involvement of break-induced-like phenomenon in the generation of DIGA.

**Figure 4.**
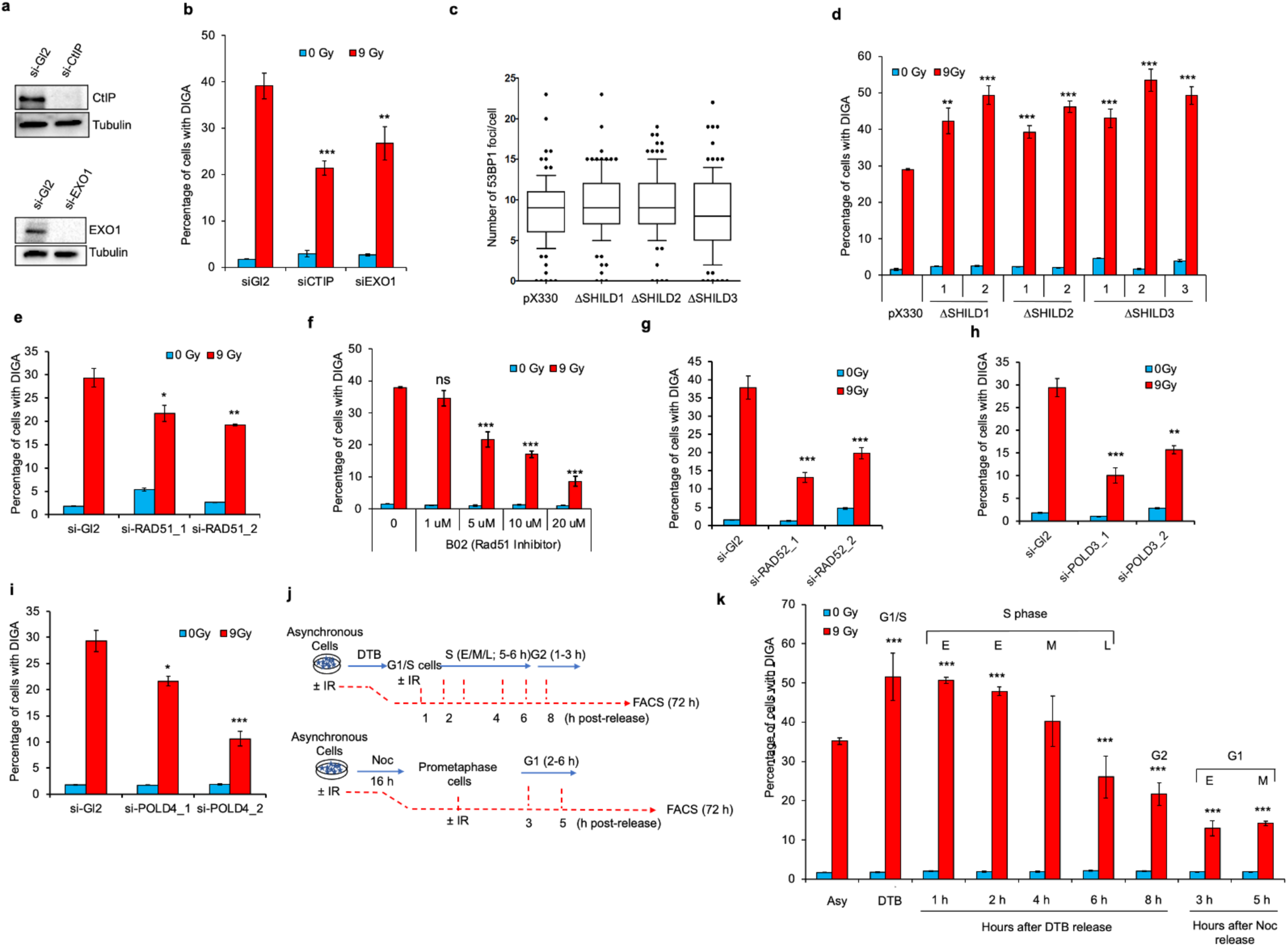
Unprotected, hyper-resected DSBs promote DIGA in mammalian cells through BIR-like mechanism. **a**, **b**, Histogram (***b***) showing the impact of IR (9 Gy) on the induction of genomic amplifications in control U2OS cells or in U2OS cells depleted of CtIP or EXO1 by siRNA (immunoblot shown in (**a**)). **c**, Quantitation of the number of 53BP1 foci per cell in control (pX330) cells or U2OS deleted of SHILD1, SHILD2, or SHILD3. Representative images are shown in the Extended Data Fig. 9a. The results represent the average of a minimum of 100 nuclei in each of three independent experiments for each condition ± S.D. ns: non-significant. **d**, Histogram showing the percentage of cells with genomic amplifications following the exposure of control (pX330) U2OS cells or individual independent clones of U2OS cells lacking SHILD1, SHILD2, or SHILD3 to 9 Gy. Genomic amplifications were monitored 72 h post IR. Representative FACS profiles shown in Extended Data Fig. 9b. **e**, Induction of genomic amplifications by IR requires RAD51. Representative FACS (PI) profiles of control U2OS cells or U2OS cells depleted of RAD51 by two different siRNAs and exposed to 9 Gy. Genomic amplifications were monitored 72 h post IR. **f**, Dose-dependent suppression of strand invasion following incubation of U2OS cells with the RAD51 recombinase inhibitor B02. Representative FACS profiles shown in Extended Data Fig. 9c. **g-i**, Induction of genomic amplifications by IR requires the break-induced replication factors RAD52, POLD3 and POLD4. Representative FACS (PI) profiles of control U2OS cells or U2OS cells depleted of RAD52 (**g**), POLD3 (**h**) or POLD4 (**i**) by two different siRNAs each and exposed to 9 Gy. Genomic amplifications were monitored 72 h post IR. **j**, **k**, Histogram (**k**) showing the percentage of U2OS cells undergoing genomic amplifications when analysed at 72 h following the exposure of U2OS synchronized at various stages of the cell cycle to 9 Gy. Cells were irradiated at Early (E), Mid (M) or late (L) S-S-phase of the cell cycle following the synchronization by DTB and release (depicted in the schematic in (**j**; top) or in early (E) or Mid (M) G1 phase of the cell cycle following the synchronization in prometaphase by nocodazole and release (depicted in the schematic in (**j**; bottom)). Genomic amplifications in all cases was monitored 72 h following the exposure to IR. The synchronization PI-FACS profiles and representative rereplication PI-FACS profiles are shown in Extended data Fig. 10. **h**, Data in all the histograms represent the average of three independent experiments ± S.D. \**p* < 0.05, \*\**p* < 0.01, \*\*\**p* < 0.001.

Since hyper-resection promotes HR when an homologous template is available (as in late S-phase or G2), and is antagonized by 53BP1-RIF1-Shieldin, our results suggest that cells encountering DSBs in late S-phase or in G2 will be less prone to DIGA induction than cells in early or mid-S-phase. To test this hypothesis, U2OS cells were synchronized at the G1/S transition by DTB and exposed to 9 Gy at various stages of the cell cycle following release from the DTB and DIGA was measured 72 h post-IR (Fig. 4j – top and Extended Data Fig. 10a). Consistent with our hypothesis, cells encountering DSBs in late S-phase or in G2 were significantly less prone to DIGA induction than cells in early/mid-S-phase (Fig. 4j-top; 4k and Extended Data Fig. 1a, b). The exposure of U2OS cells in early or mid G1 phase (following their release from the nocodazole) to IR resulted in the least DIGA induction (Fig. 4j-bottom; 4k and Extended Data Fig. 10c, d).

Our data reveal a previously unrecognized mechanism by which DSBs, induced by IR or by other clinically relevant chemotherapeutics, result in repair-associated and break-induced cytotoxic DNA synthesis. This appears to be templated by unshielded, hyper-resected ends of DSBs (potentially within replication bubbles) invading non-homologous sequences, thereby enabling long-range DNA amplifications. We speculate that sub-lethal DIGA levels may contribute to genomic amplifications and genomic instability appearing in IR-induced malignancies.

## Materials and Methods

### Cell culture

U2OS osteosarcoma cells, HEK 293T embryonic kidney cells, and breast cancer cells (MDA-MB-231 were maintained in Dulbecco’s modified Eagle’s medium (DMEM) supplemented with 10% fetal bovine serum (FBS) and 1% penicillin/streptomycin (P/S). The previously described A*si*SI-ER-U2OS cell line^22^ was maintained in the same culture media used for parental U2OS cells. The human melanoma cell lines VMM39, VMM1, VMM18, VMM5A, VMM15, and VMM12 were established from metastatic lesions of patients at the University of Virginia (IRB #5202, Dr. CL. Slingluff). DM93, DM331, DM13, DM112, DM6, and SLM2 melanoma cell lines were established from metastatic lesions by Dr. H.F. Seigler at Duke University^55–61^. SK-MEL-2 and SK-MEL-28 melanoma cells were obtained from the American Type Culture Collection (ATCC, Manassas, VA). All melanoma cells were grown in RPMI medium supplemented with 10% FBS and 1% P/S. Non-malignant human PIG3V melanocytes^62^ were maintained in Media 254 (ThermoFisher Scientific) containing 1% human melanocyte growth supplement (HMGS), 5% FBS, and 1% P/S. The human head and neck squamous cell carcinoma (HNSCC) cell lines Cal27, FaDu, and SCC25 were obtained from ATCC (Manassas, Virginia). The UNC7 HNSCC cells were provided by Dr. Wendell Yarbrough (Vanderbilt University, Nashville, Tennessee). HNSCC cells were maintained in DMEM/Ham’s nutrient mixture F12 (Sigma Aldrich) supplemented with 10% FBS and 1% P/S. Human h-Tert immortalized keratinocytes (OKF6-TERT2) were obtained from Dr. James Rhienwald (Harvard Medical School), and were cultured in GIBCO keratinocyte serum-free medium (K-sfm) supplemented with 25 μg ml^-1^ BPE, 0.4 mM CaCl2, 0.2 ng ml^-1^ EGF, and 1% P/S. The colorectal cancer HCT116 cell lines were obtained from Dr. Anindya Dutta (University of Virginia), and were maintained in McCoy’s 5A (Modified) Medium supplemented with 10% FBS and 1% P/S. All cells were grown at 37°C in 5% CO2, and were periodically tested for mycoplasma contamination (MycoAlert; Lonzo Group Ltd). The identity of the cells was validated by STR profiling (ATCC), by morphological examination, and in some cases, by analysis of chromosome number in metaphase spreads.

### Flow cytometry

The effect of IR and MLN4924 on the cell cycle and rereplication induction was assessed by flow cytometry with propidium iodide (PI) or bromodeoxy uridine (BrdU) staining. Cells were treated with various doses of MLN4924 (Active Biochem) or exposed to various doses of IR for different time points before harvesting. Cells were washed with ice-cold PBS and resuspended in ethanol (70%). Cells were subsequently treated with a PI staining buffer (50 μg ml^-1^ PI (Sigma Aldrich), 10 μg ml^-1^ DNase-free RNase A, 0.05% NP40). A FACscan (Becton Dickinson) flow cytometer was used to analyse the samples, and G0/G1, S, and G2/M fractions were segmented. Subsequent analysis using FlowJo and ModFit software was used to determine the fraction of cells with genomic content of greater than G2/M (rereplication). For BrdU staining, cells were pulse-treated with BrdU (10 nM) for 1 h in the dark before harvesting. Cells were washed with PBS and staining solution before they were fixed and permeabilized according to the manufacturer’s instructions (BD Biosciences, San Diego, CA). Cells were subsequently stained with anti-BrdU antibody solution at room temperature for 20 min, washed, and stained with 7-AAD (BD Biosciences) for 30 min at 4°C. The cells were resuspended in 1 ml of staining buffer, and stored at 4°C overnight before analysis. Samples were analysed on a FACscan (Becton Dickinson) flow cytometer, and the fraction of BrdU positive cells was determined using FlowJo and ModFit software. Where indicated, cells were treated with ATM kinase inhibitor KU-55933, DNA-PKcs kinase inhibitor NU-7441, or Rad51 inhibitor B02 (all purchased from TOCRIS Bioscience) 24 h before irradiation. The DNA LIG4 inhibitor SCR7 (#SML1546, Sigma Aldrich) was added 24 h before irradiation, and was replenished every 24 h. Analysis of rereplication induction by FACS (PI staining) following the induction of DNA damage by etoposide (1 μg ml^-1^, TOCRIS), doxorubicin (0.1 μM, Sigma Aldrich), or by UV (100 J m^2-^) was performed 72 following treatment.

### Cell synchronization and exposure to IR

Asynchronously growing U2OS cells were synchronized at the G1/S boundary by double thymidine block (DTB) and release. Briefly, U2OS cells were treated with 2 mM thymidine (ACROS Organics) for 14 h, then washed with PBS, and fresh medium was added. After 9 h, cells were treated again with thymidine (2 mM), and incubated for another 14 h. Cells were released from G1/S arrest by washing with PBS and addition of fresh media. Cell synchronization was monitored by flow cytometry. Cells synchronized at various stages of the cell cycle were exposed to 9 Gy of X-ray (Gamma irradiation was performed at the University of Virginia Small Animal Radiation Research Platform [SARRP; Xstrahl]), and were incubated for 48 or 72 h prior to analysis by FACS. In another scheme, U2OS cells were synchronized in prometaphase by treating the cells with nocodazole (0.5 μg ml^-1^, M1404; Sigma Aldrich) for 24 h. Prometaphase cells were collected by mitotic shake off, counted, washed, and re-seeded with or without nocodazole, and exposed to 9 Gy at various time points following release from the nocodazole block. Parallel samples were collected for flow cytometric analysis to determine the phase in which the various cells were exposed to IR.

### Meselson-Stahl Caesium Chloride Gradient Ultracentrifugation

U2OS cells were synchronized at the G1/S boundary using double thymidine block (DTB). BrdU (100 μM) was added upon release. Eight hours post-release from the DTB, cells were either mock-treated or irradiated with 9 Gy, and were treated with nocodazole (0.5 μg ml^-1^) and BrdU (100 μM) with or without aphidicolin (2 μM; EMD Millipore Corp). Cells were collected 36 h post-irradiation. A sample not treated with BrdU was collected in parallel, and served as an unstained negative control. Cells were lysed in lysis buffer (100 mM NaCl, 10 mM Tris-HCl pH 8, 25 mM EDTA, 0.5% SDS), and proteins were digested with 20 μg of proteinase K overnight at 55°C. Genomic DNA was extracted by Phenol/Chloroform/isoamyl alcohol extraction and ethanol precipitation. Purified DNA (50 μg in 200 μl total volume) was sheared by sonication for 2 sec into fragments ranging from 0.5 to 3 kb. Digested DNA was mixed with 0.96 g ml^-1^ Caesium Chloride in TE Buffer (giving a density of 1.71) in 13 ml Polyallomer Quick-Seal ultracentrifuge tubes (Beckman). Samples were spun at 44,000 g in a 70.1 Ti Rotor (Beckman) for 66 h. fractions (200 μl) were collected from the bottom of the tubes, and the DNA concentration was measured using Qubit fluorometric quantification. The percentage of DNA exhibiting conservative replication (containing H:H DNA) was calculated by measuring the area under the curves from three independent experiments using GraphPad Prism v.7.0.

### Detection of BrdU in CsCl fractions via ELISA

CytoSelect BrdU Competitive ELISA kit (CBA-5098; Cell Biolabs, Inc) was used to confirm BrdU incorporation in the H:H and H:L fractions collected after Caesium Chloride gradient ultracentrifugation. DNA sample preparation and plate coating were performed according to the manufacturer’s protocol. Briefly, 50 ng of DNA from selected fractions was heat-denatured to single-stranded DNA (ssDNA) and then digested into nucleosides with 20 units of Nuclease P1 (#M0660, NEB) for 2 h followed by 1 h incubation with 10 units of CIP Alkaline phosphatase (#M0290, NEB). The digested DNA samples were purified and added to a BrdU conjugate preadsorbed microplate prepared according to the manufacturer’s instructions. Anti-BrdU monoclonal antibody was added to each well and incubated for 1 h at room temperature. Following a series of three washes, an HRP-conjugated secondary antibody was added to all wells and incubated for 1 h. After 3 washes and treatment with the provided substrate/stop solutions, absorbance of each microwell was read on a plate reader at 450 nm (Synergy HT, BioTek). BrdU content was determined by comparing the absorbance values with a predetermined BrdU standard curve.

### Cell lysis, SDS-PAGE, and immunoblotting

Cells were lysed with Radioimmunoprecipitation assay (RIPA) lysis buffer (50 mM Tris, pH 8.0; 150 mM NaCl, 1% NP-40; 0.5% sodium deoxycholate; 0.1% SDS) supplemented with 1X Halt Protease and Phosphatase Inhibitor Cocktail (Thermo-Scientific). Equal amounts of protein lysates were separated on polyacrylamide 8–12% gels (BioRad, Hercules, CA) by electrophoresis and transferred to nitrocellulose membranes. Membranes were blocked with 1X phosphate buffered saline, 1% Tween 20 (Sigma Aldrich; PBST) with 5% milk (or 5% BSA) for one hour at room temperature, and incubated with the appropriate primary antibodies in 1X PBST for 1 h at room temperature or overnight at 4°C. The membranes were washed 5 times with 1X PBST and incubated with horseradish peroxidase conjugated secondary anti-mouse or anti-rabbit IgG (1:5000, Cat # P0161 and P0448, respectively; DAKO) in 1X PBST for 1 h at room temperature. Immunoblot signals were detected by enhanced chemiluminescence (Millipore). Primary antibodies recognizing the following proteins were used: ATM (1:1000, ab78; Abcam), tubulin (1:1000, sc-53646; Santa Cruz), SET8 (1:1000, C18B7; Cell Signaling), mono-methyl Histone H4 (K20) (1:500, #9724; Cell Signaling), di-methyl Histone H4 (K20) (1:500, #9759; Cell Signaling), Tri-methyl Histone H4 (K20) (1:500, #5737; Cell Signaling), DNA-PKcs (1:1000, ab44815; Abcam), 53BP1 (1:5000, N100-304; Novus Biologicals) POLD3 (1:1000, H00010714-M01; Abnova), POLQ (1:1000, H00010721-M09; Abnova), RIF1 (1:1000, A300-569A-M; Bethyl Laboratories), RNF8 (1:1000, sc-2711462; Santa Cruz), RNF168 (1:1000, ABE367; Millipore), CtIP (1:1000, sc-271339, Santa Cruz), EXO1 (1:1000, ab95012; Abcam), XRCC4 (1:1000, sc-271087; Santa Cruz), XLF (1:1000, A300-730A-M; Bethyl Laboratories), NBS1 (1:1000, ab32074; Abcam), MDC1 (1:25,000, ab11171; Abcam), LIG1 (1:1000, 18051-1-AP; Proteintech), and LIG4 (1:1000, HPA001334; Sigma Aldrich).

### Immunofluorescence

Slides were fixed in 4% paraformaldehyde (in PBS) for 10 min (or 2% for 30 min). Slides were washed three times with PBS, then permeabilized in extraction buffer (0.1% Triton-X in PBS) for 10 minutes. Slides were washed 3X in wash buffer (0.5% BSA in 1X PBS) for 5 min, blocked in 2% BSA in PBS for 45 min, washed (3x 5 min each), and then incubated with anti-53BP1 (1:1000, N100-304; Novus Biologicals) in 0.5% BSA with 0.1% Triton-X overnight at 4°C. Slides were washed and incubated with Alexa Fluor 488 anti-IgG (1:500; Life Technologies) in 0.5% BSA in PBS for 1 h at room temperature. Slides were washed, dried, and mounted with 4′, 6′-diamidino-2-phenylindole (DAPI; #H-1200, Vector Laboratories, Inc.), and 53BP1 foci were visualized and quantified using EVOS FL cell imaging system fluorescence microscope (Advanced Microscopy Group). A minimum of 100 nuclei from random fields were examined for each group.

### RNA interference

Transfections with siRNAs were performed using Lipofectamine RNAi-max according to the manufacturer’s protocol (Invitrogen, Carlsbad, CA). Cells were seeded at 25% confluence and transfected with the individual siRNAs (10 nM each) in the appropriate growth media. Cells were harvested 48 and 96 h post-transfection for cell cycle analysis. The following siRNAs were used (sense strand): si-GL2 (control): 5′-AACGUACGCGGAAUACUUCGA-3′; si-CDT1: 5′-AACGUGGAUGAAGUACCCGAC-3′; si-SET8: 5′-GAUUGAAAGUGGGAAGGAA-3′; si-EXO1: 5’-CAAGCCUAUUCUCGUAUUU-3’; si-CtIP 5’-GCUAAAACAGGAACGAAUC-3’.

### Gene targeting by CRISPR/Cas9, and establishment of individual knockout cell lines

Two single guide-RNAs (sgRNAs) targeting the various genes at two proximal (within 300 bp) sites in early exons of each gene were cloned into pX330 vector (Addgene #42230) containing a human codon-optimized *Sp*Cas9 endonuclease using BbsI restriction enzyme to cut sites downstream of the U6 promoter. Plasmids were amplified in DH5α bacteria (Invitrogen), purified using the QIAprep Spin Miniprep Kit according to the manufacturer’s instructions (Qiagen), and verified by Sanger sequencing (Eurofins Scientific) using U6-specific primers. Various cell lines were transfected along with pMSCV vector containing the puromycin resistance gene (Clontech) using Lipofectamine 2000 according to the manufacturer’s instructions (Invitrogen). At 24 h post-transfection, cells were incubated with puromycin (2 μg ml^-1^) for 48 h, and were subsequently seeded at low density to obtain single colonies. Individual clones were propagated in culture in the appropriate growth medium, and aliquots were frozen. Samples of each clone were lysed overnight at 55°C in lysis buffer (100 mM NaCl, 10 mM Tris-HCl pH 8, 25 mM EDTA, 0.5% SDS) supplemented with 20 μg proteinase K. Genomic DNA was extracted by phenol/chloroform/isoamyl alcohol followed by ethanol precipitation. Genotyping was performed using PCR amplification of the targeted locus with gene-specific primers followed by Sanger sequencing (Eurofins Scientific). The deletion of the various genes in the knockout-positive cell lines was further confirmed by immunoblotting. The following sg-RNAs (sense strand) were used to generate the various knockout sgRNA plasmids in the pX330 backbone: ATM: sg-ATM-1: 5’-TCAACTAGAACATGATAGAG-3’, sg-ATM-2: 5’-GATTCGAGATCCTGAAACAA-3’; PRKDC: sg-DNA-PKcs-1: 5’-GAGCCGGTGTGCGTTGCTCC-3’, sg-DNA-PKcs-2: 5’-GCCGGTCATCAACTGATCCG-3’; TP53BP1: sg-53BP1-1: 5’-GACGCACAAAGAAAATCCTG-3’, sg-53BP1-2: 5’-GAACGAGGAGACGGTAATAG-3’; XRCC4: sg-XRCC4-1: 5’-AGTATAACTCATTTTCTACA-3’, sg-XRCC4-2: 5’-TTTGTTATTACACTTACTGA-3’; RIF1: sg-RIF1-1: 5’-CCTCGCGCCGCTGTTGGAGA-3’, sg-RIF1-2: 5’-ACGCTTACCTGACTCTGACC-3’; RNF8: sg-RNF8-1: 5’-CCGGGGTCGAGTAGGCGATG-3’, sg-RNF8-2: 5’-TTCGTCACAGGAGACCGCGC-3’; RNF168: sg-RNF168-1: 5’-TCGCTGTCCGAGTGCCAGTG-3’, sg-RNF168-2: 5’-GGTATCGTCGTGGACTCGGT-3’; NHEJ1: sg-XLF-1: 5’-TGGGCGTGGCTACAGCTTGC-3’, sg-XLF-2: 5’-TGAACAGGTGGACACTAGTG-3’; MDC1: sg-MDC1-1: 5’-GGACACCCAGGCTATTGACT-3’, sg-MDC1-2: 5’-GTAGGGCGGCTACATATCTT-3’; LIG1: sg-LIG1-1: 5’-AGAGTGACTCTCCGGTGAAG-3’, sg-LIG1-2: 5’-TTAGCCCTGCTAAAGGCCAG-3’; LIG4: sg-LIG4-1: 5’-CACAAACTTCACAAACTGTT-3’, sg-LIG4-2: 5’-GCAATGAGACTAATTCTTCC-3’; POLQ: sg-POLQ-1: 5’-TGAATCTTCTGCGTCGGAGT-3’, sg-POLQ-2: 5’-GATTCGTTCTCGGGAAGCGG-3’; KMT5B: sg-SUV4-20H1-1: 5’-TGACTAAATGCACCTGGGTC-3’, sg-SUV4-20H1-2: 5’-AAAATTACAGCACACGGGGA-3’; KMT5C: sg-SUV4-20H2-1: 5’-GCCGGAAAGTGGCTTTACCA-3’, sg-SUV4-20H2-2: 5’-AGATCGTGTCCACTCGTGCT-3’; SHLD1: sg-SHLD1-1: 5’-TCAGCGTGTGACATAAGAGA-3’, sg-SHLD1-2: 5’-ACAGCGAGGCTTTCAGTTCT-3’; SHLD2: sg-SHLD2-1: 5’-ATTGGTTCTCCAGATCTTAG-3’, sg-SHLD2-2: 5’-CTAGACTGAGTGATATAACT-3’; SHLD3: sg-SHLD3-1: 5’-TGTGAGAGTGATCCCACACA-3’, sg-SHLD3-2: 5’-AGCTTCCACTCAGACCTAAA-3’.

### Clonogenic survival assays

Radiation sensitivity in the various cell clones was established using a clonogenic survival assay. Briefly, the various cell lines were trypsinized, counted using a Countess Automated Cell Counter (Invitrogen), and serial dilutions of cells were seeded in 10 cm dishes in triplicate. Twenty-four hours after seeding, cells were irradiated (1-9 Gy) and cultured for two weeks. Once colonies reached the appropriate size (>50 cells each), cells were washed in ice-cold PBS, fixed in cold methanol for 10 min, and stained with crystal violet (0.5%) for 10 min. Plates were washed with water, dried, and images were captured using Imagelab software (BioRad). QuantityOne software (BioRad) was used to quantitate the number of colonies, and survival curves were established based on the linear quadratic model, using the formula S=e^−αD-βD2^; where S represents the surviving fraction and D represents the dose of irradiation. Results are represented as mean ± standard deviation (SD) from three independent experiments normalized to the corresponding non-irradiated plates for each group.

### Statistical analyses

All statistical analyses were performed using Excel and GraphPad Prism v. 7.0. At least three independent experiments were performed for each data set. No statistical methods were used to predetermine the sample size. Numerical data were expressed as mean ± standard deviation (SD) and the statistical significance in each case was calculated by two-tailed Student’s *t*-test (Mann-Whitney *U*-test). *P*-values were determined for all analyses, and *p* < 0.05 was considered significant. \**p* < 0.05, \*\**p* < 0.01, \*\*\**p* < 0.001. For correlations, a Spearman correlation was used, and *p*-values < 0.05 were considered significant.

## Acknowledgments

We would like to thank Drs. C.L. Slingluff, K. Lee, and E. Shibata for providing valuable reagents. We thank Dr. Y. Wang for critical reading of the manuscript and for valuable suggestions. We also thank the FACS core Facility for providing technical assistance, and Dr. J. Larner for valuable discussions. This study was supported by GM135376 from the National Institute of General Medical Sciences (NIGMS) of the NIH (T.A.).

## Author Contributions

M.B., R.E., K.D., and T.A. designed and performed experiments, and interpreted results. T.A. wrote the manuscript with input from all authors.

---

**Extended Data Figure 1:**
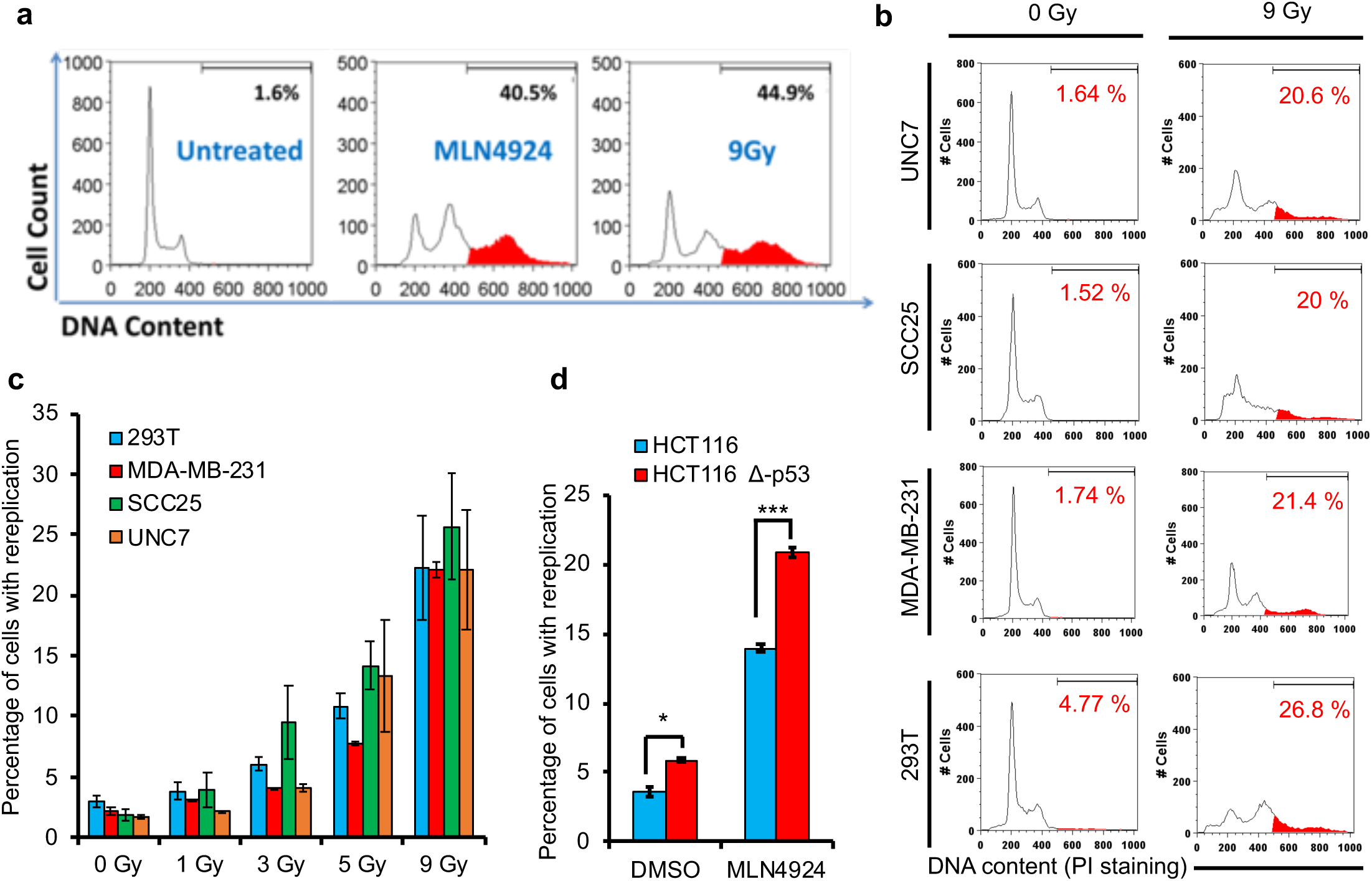
Induction of genomic amplifications in cancer cells by ionizing radiation. **a**, Representative flow cytometry (FACS) profiles (PI staining) showing the increase in genomic content greater than G2/M (highlighted in red) in FaDu cells exposed to 0.1 μM MLN4924 or 9 Gy, and examined 72 h post-treatment. **b**, Representative flow cytometry profiles of the indicated cancer cell lines showing the extent of rereplication as determined by PI-FACS 48 h following exposure to 9 Gy. **c**, Histogram showing the percentage of the indicated cancer cell lines with genomic amplifications 48 h following exposure to increasing doses of IR. Data represent the average of three independent experiments ± S.D. **d**, The impact of p53 on the percentage of HCT116 colon cancer cells exhibiting rereplication 48 h after treatment with 0.3 μM MLN4924. Data represent the average of three independent experiments ± S.D. \**p* < 0.05, \*\*\**p* < 0.001.

**Extended Data Figure 2:**
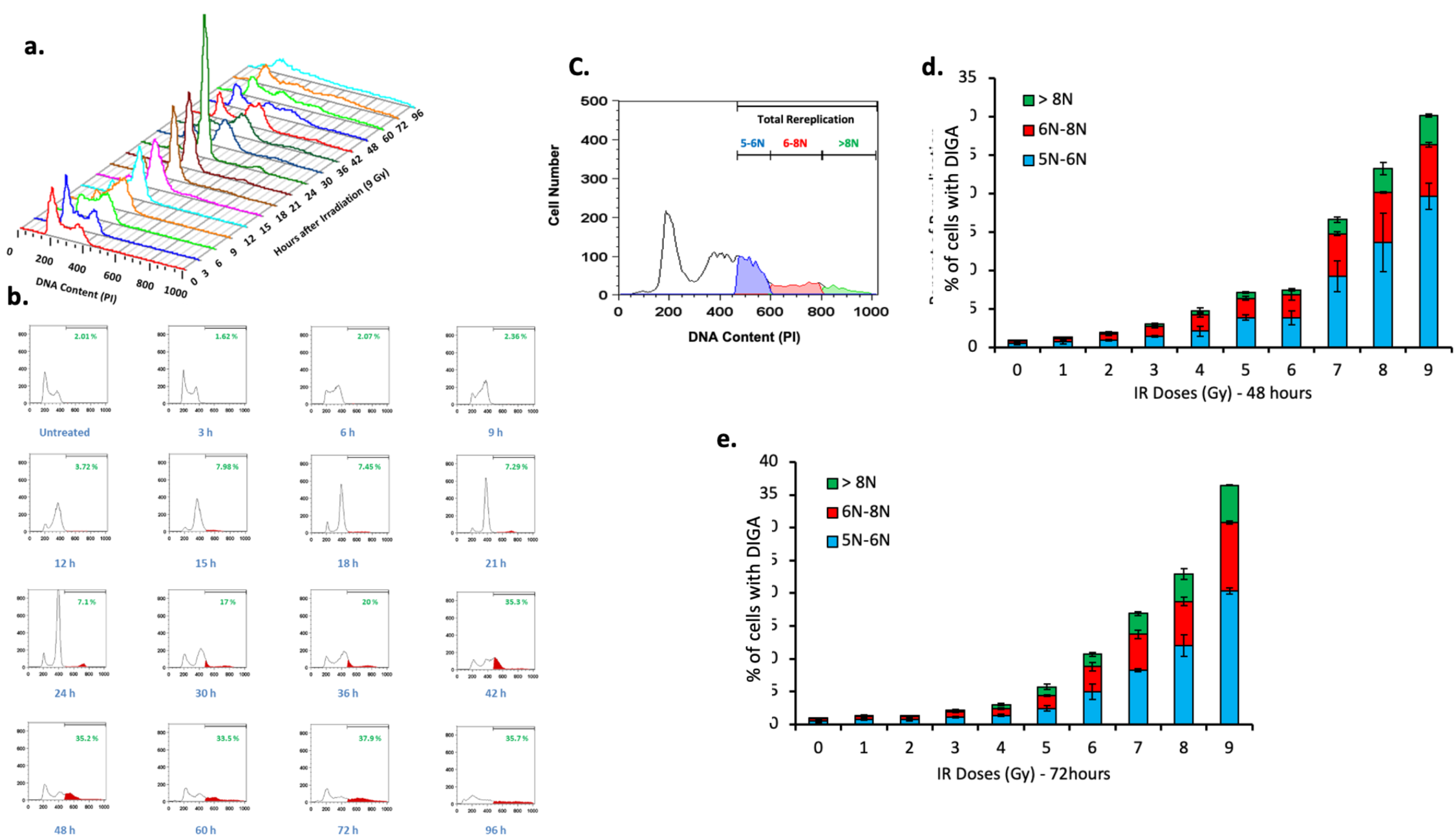
Time-dependent accumulation of heterogenous (sizes) genomic amplifications following the exposure to IR. **a**, time-dependent increases in genomic amplifications following the exposure of U2OS cells to 9 Gy. Top Flow cytometry profile alignment to show the impact of IR on the increase of >G2/M DNA content as a function of time. **b**, Representative flow cytometry profile showing the impact of IR (9G) on cell cycle distribution as a function of time, as indicated. Areas of genomic amplifications (with Percentage) are shown in red. c, Flow cytometry profile of U2OS depicting the gradual increase in DNA copy number (as measured by increased fluorescence (showing divisions to be used as cut offs for quantitation of copy # size in **d** and **e**) following the exposure of U2OS to 9 Gy. **d**, **e**, Histograms showing the increase in genomic amplifications (stratified by size in **c**) in U2OS cells exposed to increasing doses of IR (as indicated) and measured 48 (**d**) and 72 (**e**) hours post-IR exposure. Data represent the average of three independent experiments ± S.D. \**p* < 0.05, \*\*\**p* < 0.001.

**Extended Data Figure 3:**
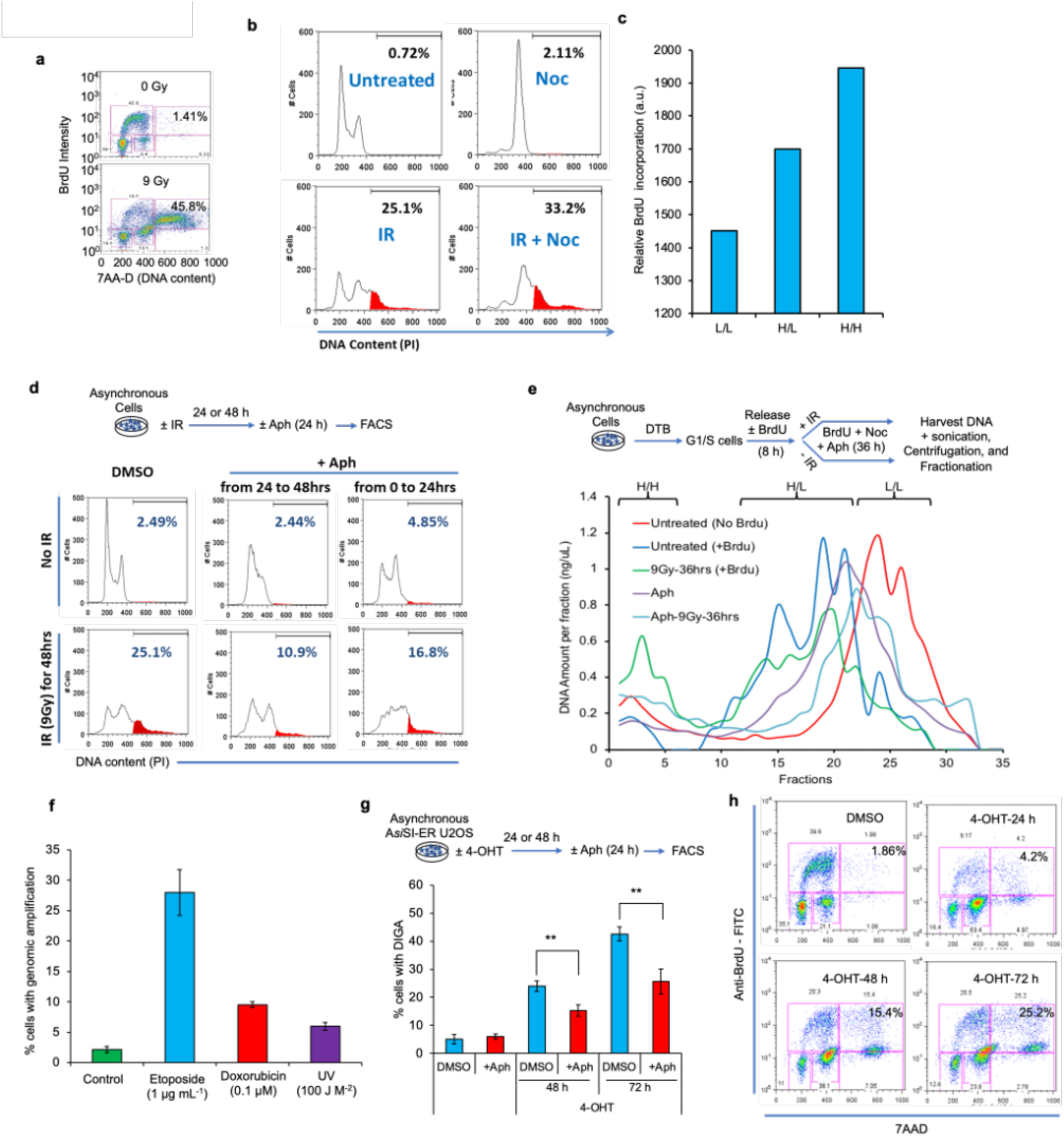
DNA double strand breaks (DSBs) induce *bona fide* genomic amplifications in cancer cells. **a**, FACS profiles showing *de novo* DNA synthesis (BrdU incorporation) in U2OS cells with greater than G2/M DNA content (as assessed by 7AAD staining) 48 or 72 h following exposure to 9 Gy. Cells were pulsed with BrdU for 1 h prior to harvest. **b**, Representative FACS profiles (PI staining) of U2OS cells 48 h following exposure to 9 Gy with or without nocodazole (Noc) treatment at the 24-48 h time point following IR exposure. Cells with genomic amplifications highlighted in red. Quantitation is shown in Fig. 1f. **c**, Elisa-based quantitation of BrdU incorporation in the peak fractions of DNA (50 ng each of L:L, H:L, and H:H DNA) isolated from the CsCl ultracentrifugation gradient shown in Fig. 1h. **d**, Workflow (*top*) and representative FACS profiles (PI staining; *bottom*) of U2OS cells left untreated or exposed to IR for 48 h and treated with or without aphidicolin (Aph) added immediately following IR and washed 24 h after (0-24 h) or added 24 h post-IR (24-48 h). Cells with genomic amplifications are highlighted in red, and quantitation of the results is shown in Fig. 1j. **e**, The appearance of H:H DNA in cells exposed to IR is aphidicolin-sensitive. The line histogram is an extension of the plot shown in Fig. 1h, and shows additional treatment with aphidicolin (Aph) with or without IR treatment as depicted in the experimental workflow shown on top. **f**, Histogram showing the percentage of U2OS cells with genomic amplifications (as determined by PI-FACS) 72 h following treatment with etoposide (1 μg ml^-1^), doxorubicin (0.1 μM), or exposure to ultraviolet radiation (UV; 100 J m^-2^). Data represent the average of three independent experiments ± S.D. **g**, Work flow (*top*) and quantitation of the percentage of A*si*SI-ER-U2OS cells with genomic amplifications (as determined by PI-FACS) following the induction of DSBs by the addition of 4-OHT for 48 h. Cells were treated with or without aphidicolin (Aph) added together with 4-OHT and washed 24 h after (0-24 h), or with Aph added 24 or 48 h following treatment with 4-OHT and harvested 24 h after Aph (48 and 72 h following treatment with 4-OHT, respectively). Data represent the average of three independent experiments ± S.D. \*\**p* < 0.01. **h**, Representative FACS profiles showing *de novo* DNA synthesis (BrdU incorporation) in A*si*SI-ER-U2OS cells with greater than G2/M DNA content (as assessed by 7AAD staining) 48, 72, or 96 h following treatment with 300 nM 4-hydroxy-tamoxifen (4-OHT). Cells were pulsed with BrdU for 1 h prior to harvest.

**Extended Data Figure 4:**
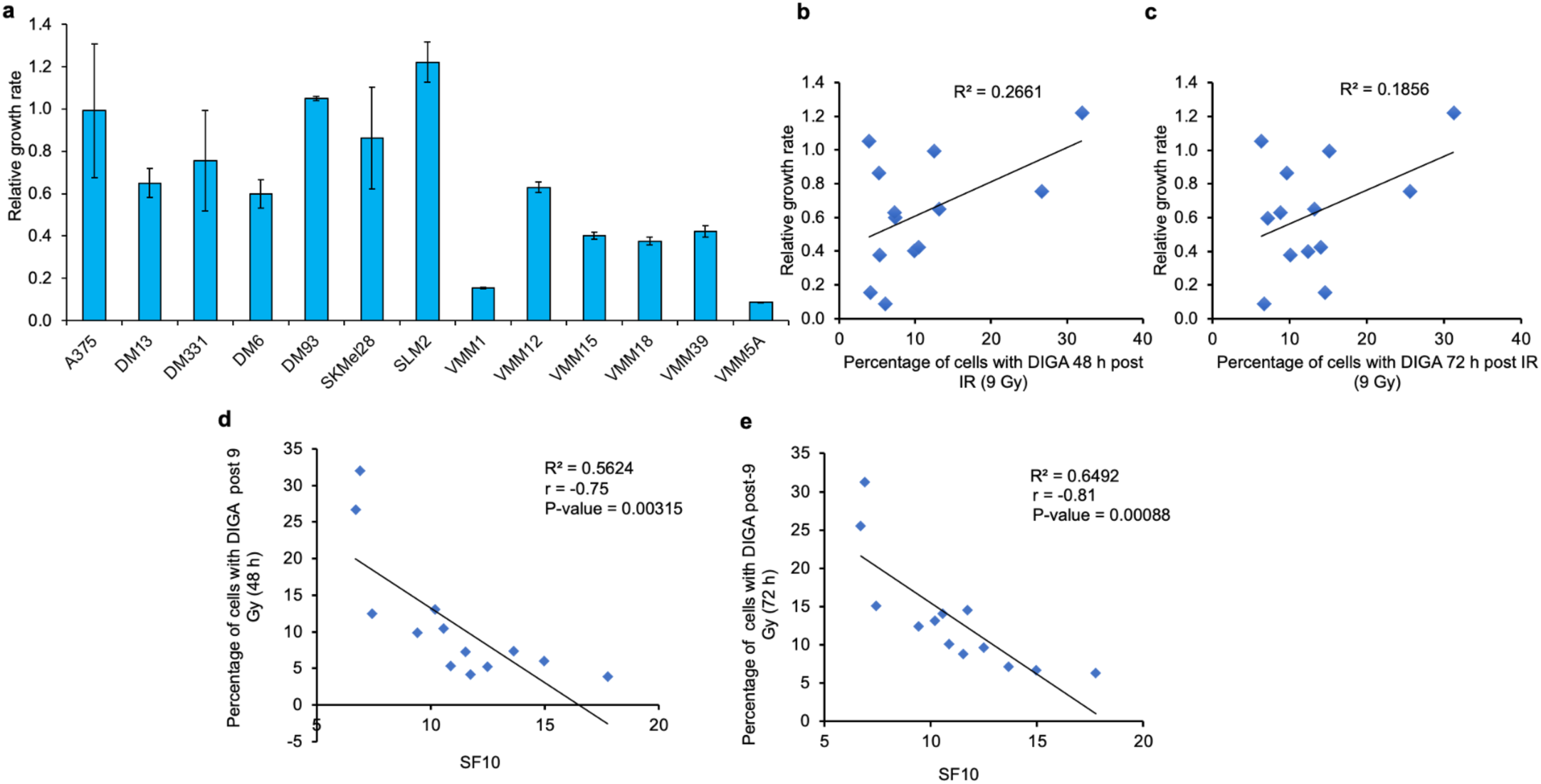
Induction of genomic amplifications in melanoma cells by IR correlates with IR-induced toxicity but not with melanoma cell proliferation. **a**, Histogram showing the relative growth rate of the indicated melanoma cells determined by cell counting. Data represent the average of three independent experiments ± S.D. **b**, **c**, Histograms showing the correlation between the percentage of melanoma cells (13 cell lines) with genomic amplifications induced 48 h (**b**) or 72 h (**c**) following exposure to 9 Gy in three independent experiments (Fig. 1l), and the doubling time of the various melanoma cells determined by cell counting in three independent growth curves. **d**, **e**, Histograms showing the correlation between the percentage of melanoma cells (13 cell lines) with the extent to which genomic amplifications are increased as determined by PI-FACS at 48 h (**d**) or 72 h (**e**) following exposure to 9 Gy in three independent experiments (shown in Fig. 1l), and the surviving fraction (at 10% of the population) determined from clonogenic survival curves in three independent experiments.

**Extended Data Figure 5:**
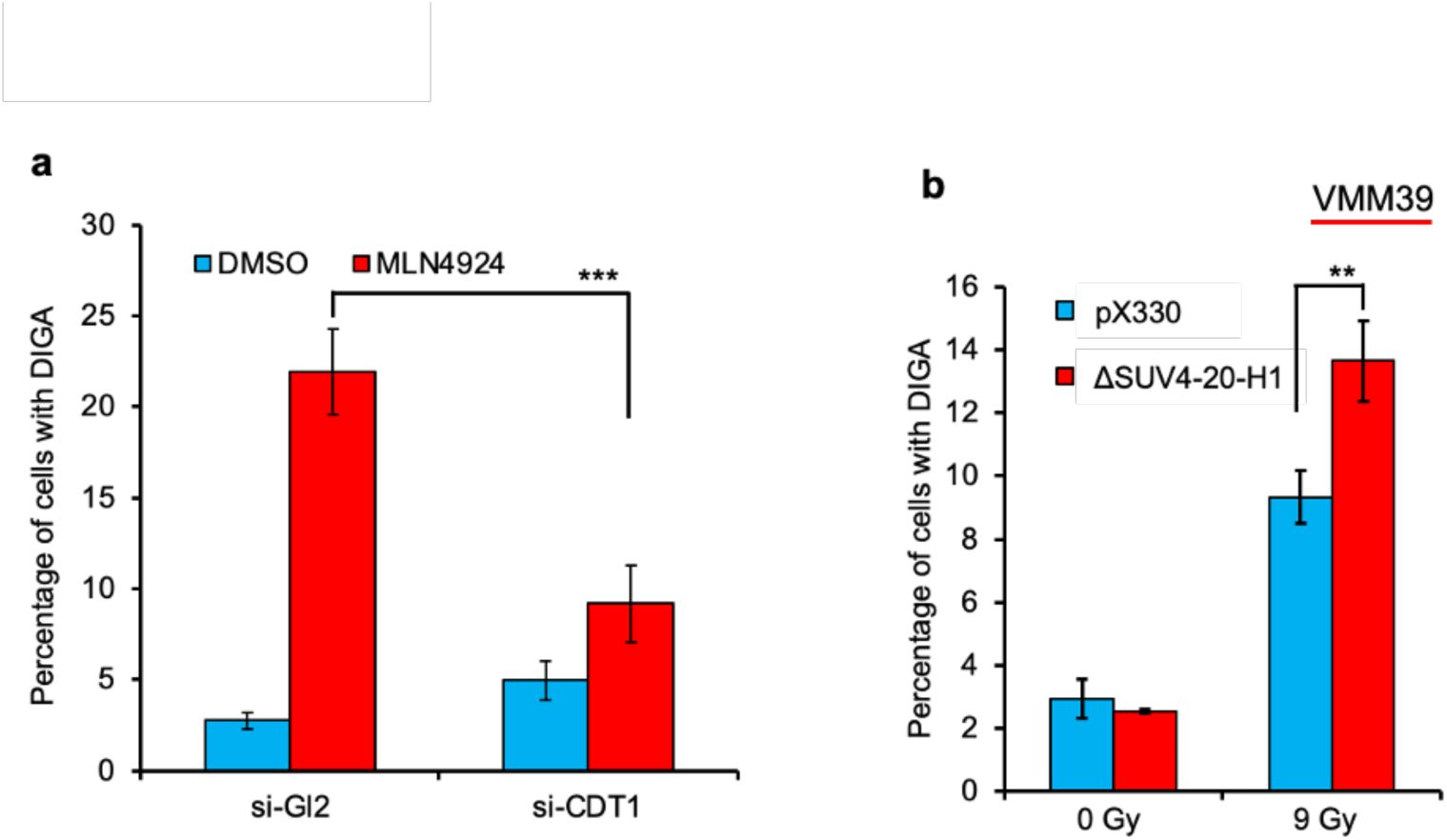
Rereplication induction by MLN4924 is CDT1-dependent, and IR-induced genomic amplifications are suppressed by SUV4-20H1. **a**, CDT1 is critical for rereplication induction by MLN4924. The Histogram shows the percentage of cells with rereplication (assessed by PI-FACS) following the treatment of control U2OS cells (si-Gl2) and U2OS cells depleted of CDT1 for 72 h prior to treatment with MLN4924 for 48 h. Data represent the average of three independent experiments ± S.D. \*\*\**p* < 0.001. **b**, SUV4-20H1 suppresses the induction of genomic amplifications in VMM39 melanoma cells. The histogram shows the percentage of control (pX330) VMM39 cells or VMM39 cells with *KMT5B* deletion with genomic amplifications determined by PI-FACS 48 h post-exposure to 9 Gy. Data represent the average of three independent experiments ± S.D. \*\**p* < 0.01.

**Extended Data Figure 6:**
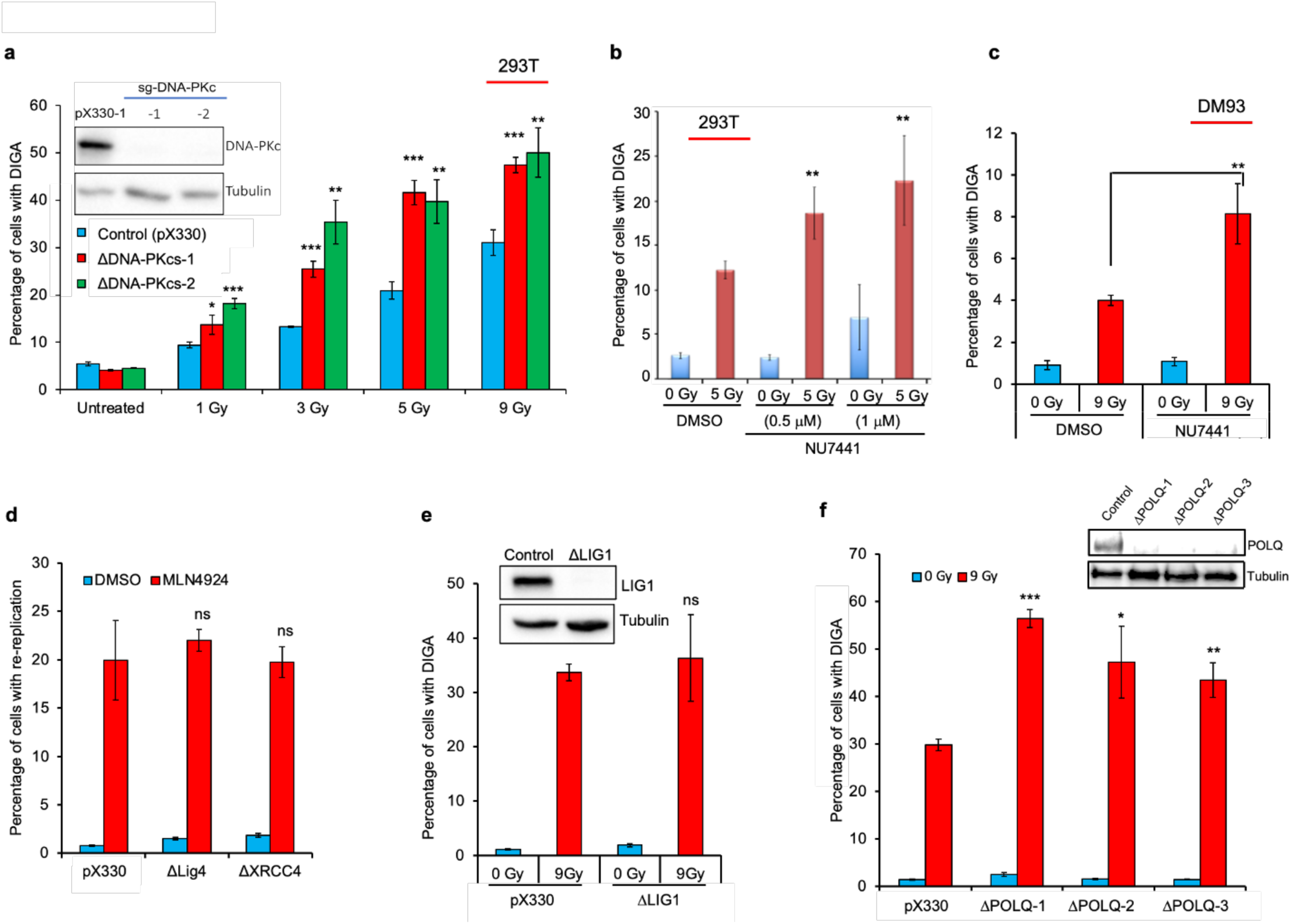
The impact of deletion or pharmacological inhibition of various proteins involved in DSB repair on the induction of genomic amplifications by IR. **a**-**c.** DNA-PKcs suppresses IR-induced genomic amplifications in cancer cells. **a**, Deletion of *PRKDC* (coding for DNA-PKcs) in 293T cells stimulates the induction of genomic amplifications by IR. The histogram shows the percentage of cells with genomic amplifications (as determined by PI-FACS) in control (pX330) 293T cells or 293T deleted of DNA-PKcs (two independent clones; shown in the immunoblot; inset) following exposure to the indicated doses of IR. Genomic amplifications were assessed 48 h post-IR. **b**, **c,** Pharmacological inhibition of DNA-PKcs stimulates the induction of genomic amplifications by IR in 293T cells (**b**) or DM93 melanoma cells (**c**). The histograms show the percentage of cells undergoing genomic amplifications (as determined by PI-FACS) 48 h (293T cells) or 72 h (DM93 cells) following exposure to 5 Gy (293T cells) or 9 Gy (DM93 cells). Where indicated, the cells were pre-treated with the DNA-PKcs specific inhibitor NU7441 for 24 h prior to IR exposure. **d**, Histogram showing the percentage of control U2OS cells (pX330) or U2OS cells deleted of LIG4 or XRCC4 (immunoblots shown in Fig. 3c) undergoing genomic amplifications (as determined by PI-FACS) 72 h following treatment with control DMSO or 0.1 μM MLN4924. **e**, Histogram showing the percentage of control U2OS cells (pX330) or U2OS cells deleted of LIG1 undergoing genomic amplifications 72 h following exposure to 9 Gy. **f**, Histogram showing the percentage of control (pX330) U2OS cells or U2OS cells deleted of POLQ (three independent clones; shown in the immunoblot; inset) undergoing genomic amplifications as determined by PI-FACS 72 h following the exposure to 9 Gy. Data in all histograms represent the average of three independent experiments ± S.D. ns: non-significant, \**p* < 0.05, \*\**p* < 0.01, \*\*\**p* < 0.001.

**Extended Data Figure 7:**
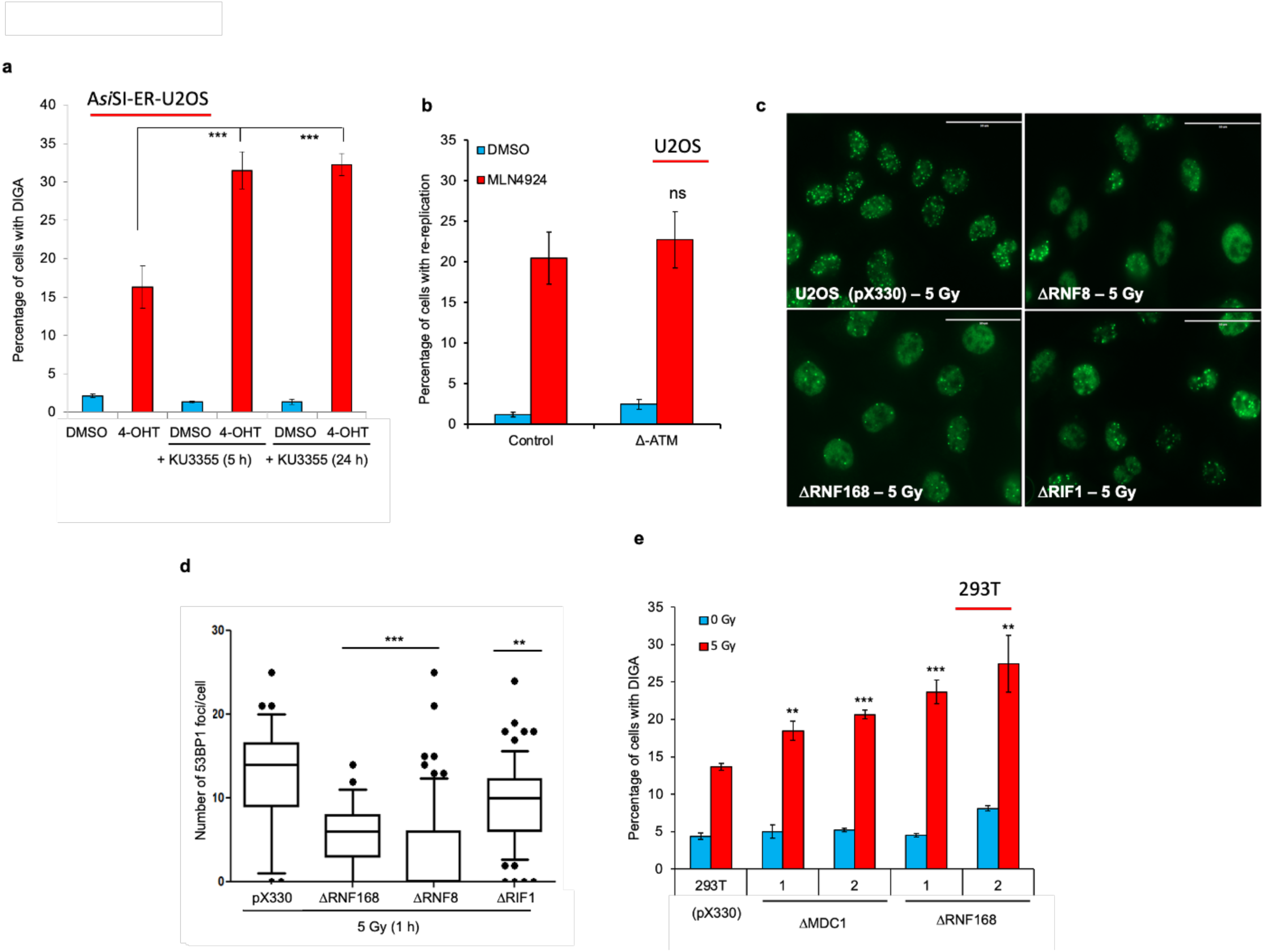
DIGA, but not rereplication induced by deregulated origin licensing, is stimulated by the deletion of factors involved in the recruitment of 53BP1 to DSB sites. **a**, ATM suppresses the induction of genomic amplifications by DSBs. The histogram shows the percentage of A*si*SI-ER-U2OS cells with genomic amplifications as determined by PI-FACS 72 h following treatment with 4-OHT. Where indicated, the cells were pre-treated with the ATM specific inhibitor KU3355 5 or 24 h prior to treatment with 4-OHT. The results represent the average of three independent experiments ± S.D. \*\*\**p* < 0.001. **b**, ATM inhibition does not stimulate rereplication induction by MLN4924. The histogram shows the percentage of control (pX330) U2OS cells or U2OS cells with ATM deletion (ΔATM) with rereplication as determined by PI-FACS 48 h following treatment with 0.1 μM MLN4924. The results represent the average of three independent experiments ± S.D. ns: non-significant. **c**, **d**, Deletion of RNF8, RNF168, or RIF1 suppresses 53BP1 recruitment to DSBs. **c,** Representative immunofluorescence images of 53BP1 showing formation of 53BP1 foci following exposure of control (pX330) U2OS cells or U2OS cells deleted of RNF8, RNF168, or RIF1 to 5 Gy and analysed 1 h post-exposure. **d**, Quantitation of the number of 53BP1 foci per cell in control (pX330) cells or U2OS deleted of RNF8, RNF168, or RIF1. The results represent the average of a minimum of 100 nuclei in each of three independent experiments for each condition ± S.D. \*\**p* < 0.01, \*\*\**p* < 0.001. **g**, MDC1 and RNF168 suppress IR-induced genomic amplifications. Histograms showing the percentage of control (pX330) 293T cells or 293T cells deleted of MDC1 or RNF168 with genomic amplifications as determined by PI-FACS 48 h following the exposure to 5 Gy. The results represent the average of three independent experiments ± S.D. \*\**p* < 0.01, \*\*\**p* < 0.001.

**Extended Data Figure 8:**
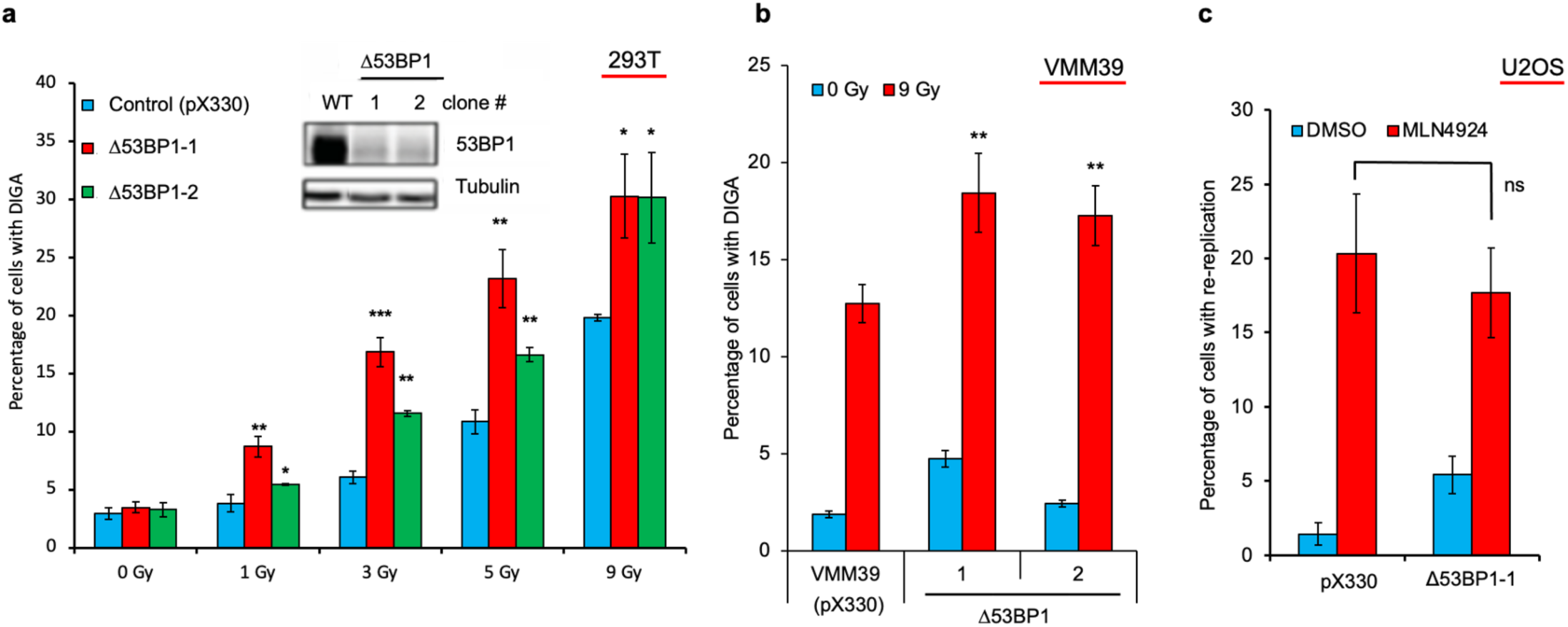
53BP1 suppresses IR-induced genomic amplifications but does not impact rereplication induction by deregulated origin licensing. **a**, Deletion *53BP1* in 293T cells enhances the induction of genomic amplifications by IR. The histogram shows the percentage of control (pX330) 293T cells or 293T deleted of 53BP1 (two independent clones; shown in the immunoblot; inset) undergoing genomic amplifications as determined by PI-FACS following exposure to the indicated doses of IR. Genomic amplifications were assessed 48 h post-IR exposure. **b**, Deletion of *53BP1* in VMM39 cells enhances the induction of genomic amplifications by IR. The histogram shows the percentage of control (pX330) VMM39 melanoma cells or VMM39 cells deleted of 53BP1 (two independent clones) undergoing genomic amplifications as determined by PI-FACS 72 h following exposure to 9 Gy. **c**, Deletion of *53BP1* does not stimulate rereplication induction by MLN4924. The histogram shows the percentage of control (pX330) U2OS cells or U2OS cells with 53BP1 deletion (Δ53BP1) with rereplication as determined by PI-FACS 48 h following treatment with 0.1 μM MLN4924. Data in all histograms represent the average of three independent experiments ± S.D. ns: non-significant, \**p* < 0.05, \*\**p* < 0.01, \*\*\**p* < 0.001.

**Extended Data Figure 9:**
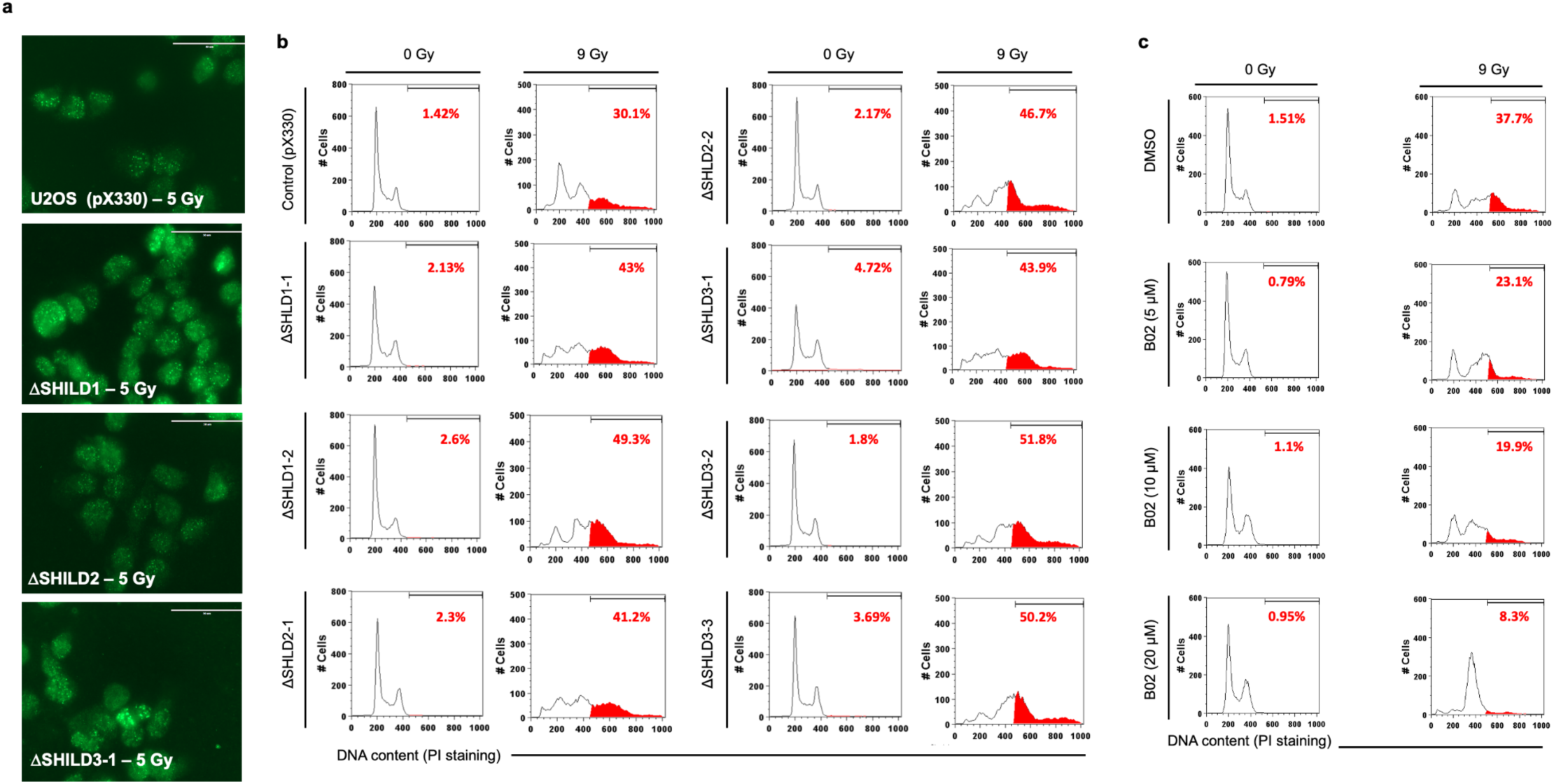
The shieldin complex and RAD51 protect cancer cells against IR-induced genomic amplifications. **a**, **b**, Deletion of SHLD1, SHLD2, or SHLD3 in U2OS cells does not impact the recruitment of 53BP1 to DSBs. **a**, Representative immunofluorescence images of 53BP1 showing formation of 53BP1 foci following exposure of control (pX330) U2OS cells or U2OS cells deleted of SHLD1, SHLD2, or SHLD3 to 5 Gy and analysed 1 h post-exposure. Quantitation of the number of 53BP1 foci per cell in control (pX330) cells or in U2OS deleted of SHLD1, SHLD2, or SHLD3 is shown in Fig. 4c. **b**, Representative flow cytometry profiles show the increase in genomic content greater than G2/M (highlighted in red) in control (pX330) U2OS cells or in cells deleted of SHLD1, SHLD2, or SHLD3 (multiple independent clones shown) exposed to 9 Gy and examined 72 h post-IR. Quantitation of the data is shown in Fig. 4d. **c**, The induction of genomic amplifications by IR requires the recombinase activity of RAD51. Representative FACS (PI) profiles of U2OS cells treated with DMSO or increasing concentrations of the RAD51 specific inhibitor B02, and exposed to 9 Gy. Genomic amplifications (highlighted in red, as determined by PI staining) were assessed 72 h post-IR exposure. Quantitation of the results is shown in Fig. 4f.

**Extended Data Figure 10:**
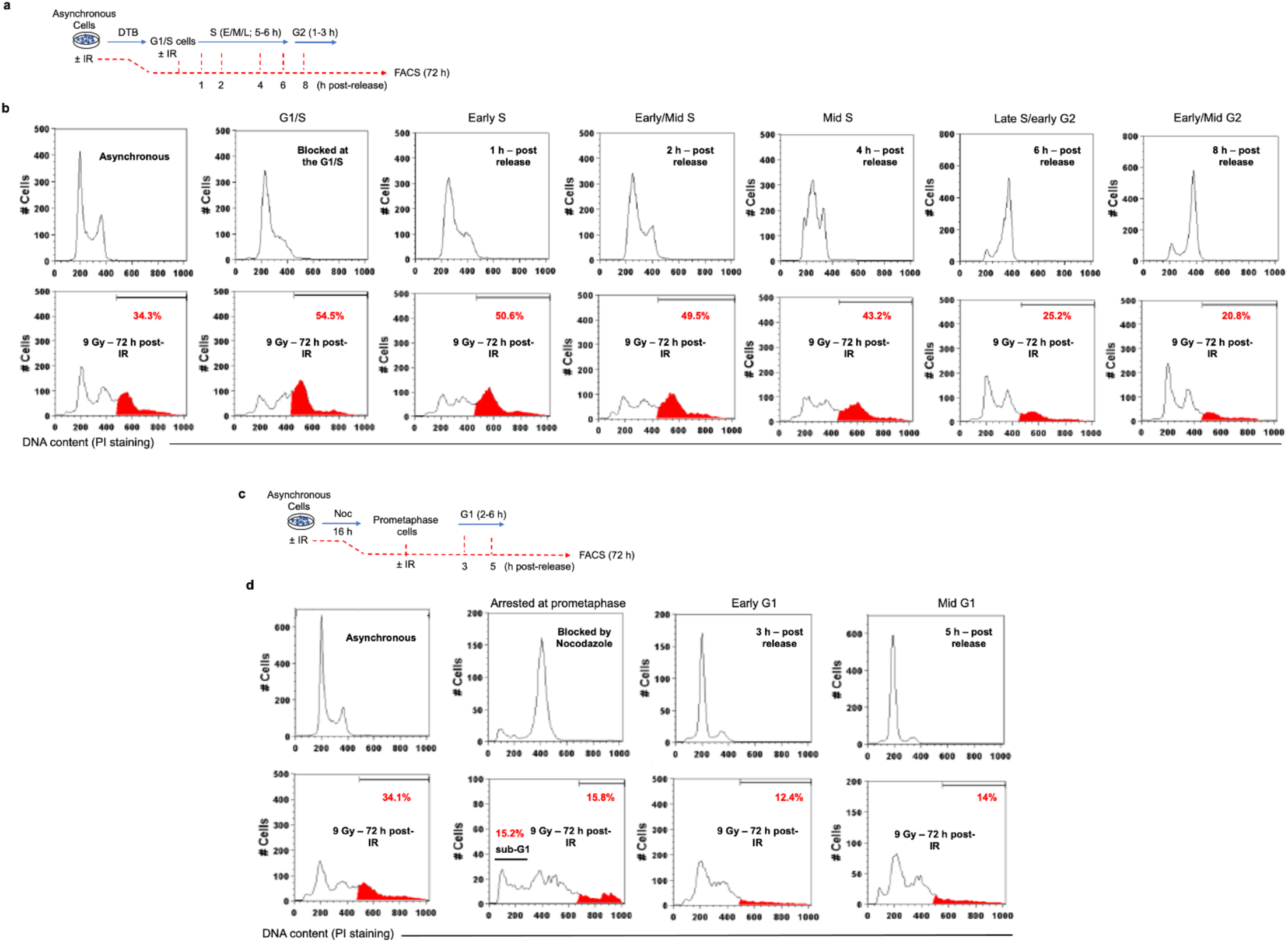
The impact of cell cycle on rereplication induction by IR. **a**, **b**, Experimental workflow (**a**) for the synchronization of U2OS cells at the G1/S by DTB and subsequent release. Cells were either harvested for PI-FACS analysis (**b**; **top panel**) at the indicated time points or exposed to 9 Gy at these time points and harvested 72 h post-IR (**b**; **bottom panel**) to monitor genomic amplifications by PI-FACS. **c**, **d**, Experimental workflow (**c**) for the synchronization of U2OS cells in prometaphase by nocodazole treatment (0.5 μg ml^-1^) and subsequent release. Cells were either harvested for PI-FACS analysis (**c**; **top panel**) at the indicated time points or exposed to 9 Gy at these time points and harvested 72 h post-IR (**d**; **bottom panel**) to monitor genomic amplifications by PI-FACS. Cells with genomic amplifications and their percentages are highlighted in red. Representative FACS profiles of each sample (in **b** and **d**) are shown. Quantitation from three independent synchronization and treatment experiments is shown in Fig. 4k.

## Notes

### Competing Interest Statement

The authors have declared no competing interest.

